# Capturing Multi-Scale Dynamics of Aortic Valve Calcification With a Coupled Fluid–Structure and Systems Biology Model

**DOI:** 10.64898/2025.11.28.691136

**Authors:** Michael Quan, Tianyou Xie, Leonard A. Harris, Haoxiang Luo

## Abstract

Calcific aortic valve disease (CAVD) arises from coupled interactions between blood flow, tissue mechanics, and cellular signaling. Hemodynamic forces influence endothelial and interstitial cell behavior, while the resulting tissue remodeling alters valve motion and flow patterns. Capturing this two-way feedback requires models that integrate fluid–structure mechanics with biochemical regulation, yet such multiscale coupling remains technically challenging. Previous computational models have focused on isolated aspects of the disease: fluid–structure interaction (FSI) simulations reproduce valve deformation and flow, and systems biology (SB) models describe molecular signaling that drives fibrosis and calcification. However, without coupling, these approaches cannot predict how mechanical dysfunction initiates biochemical remodeling or how biochemical changes feed back on mechanics. Here, we present a proof-of-principle, multi-physics computational framework that couples three-dimensional FSI simulations of aortic valve dynamics with a mechanistic SB model of calcification signaling. The FSI module resolves pulsatile blood flow and leaflet deformation, yielding local wall shear stresses and tissue strains throughout the cardiac cycle. These mechanical quantities are used as inputs to the SB module, which comprises key biochemical pathways governing inflammation, TGF-*β*/SMAD signaling, and nitric-oxide (NO)–mediated inhibition within valvular cells. Simulations predict long-term calcification trajectories for valves of varying thickness, showing that fibrosis-induced stiffening lowers shear stress, reduces NO synthesis, and enhances TGF-*β* activation, thereby accelerating calcification. While the current *one-way coupling* implementation is not intended yet for clinical applications, the framework is modular and extensible, allowing for future enhancements that will advance towards this goal. These include the incorporation of additional biological pathways in the SB model and implementation of a fully *two-way coupling* scheme between the FSI and SB models that will increase accuracy and predictive capability of the framework. By integrating physics-based hemodynamics with systems-level biochemistry, this study demonstrates the utility of a next-generation, multi-scale modeling platform for studying cardiovascular disease that unites blood flow dynamics and biochemical signaling.

## 1 Introduction

The aortic valve is a dynamic, load-bearing structure that experiences substantial mechanical stress with every cardiac cycle. Over time, this stress contributes to pathological remodeling and calcification of the valve leaflets due to a complex interplay between hemodynamic forces, cellular signaling, and tissue remodeling.^1, 2^ Progressive calcification restricts valve opening, increases left ventricular afterload, and ultimately leads to aortic stenosis and heart failure. Despite the growing prevalence of calcific aortic valve disease (CAVD), no pharmacological therapies currently exist to halt or reverse its progression, and valve replacement remains the only effective treatment.^1, 3^ Mechanobiological studies have shown that CAVD is not a passive degenerative process but a dynamic, regulated sequence of events involving endothelial activation, inflammatory infiltration, and maladaptive remodeling of the extracellular matrix.^4, 5^ Hemodynamic forces are key determinants of disease initiation and progression: the aortic and ventricular surfaces of the valve experience distinct shear stress patterns that differentially regulate nitric oxide (NO) signaling and inflammation,^6, 7^ while altered mechanical strain promotes myofibroblast differentiation and calcific nodule formation.^8, 9^ These observations indicate that valve pathology emerges from the reciprocal interplay between blood flow, tissue mechanics, and cellular signaling. Capturing this reciprocity requires models that account for both the mechanical environment that acts on valve cells and the intracellular pathways that determine how those cells respond.

A substantial body of computational research has been devoted to modeling the mechanics and hemodynamics of the aortic valve. Three-dimensional fluid–structure interaction (FSI) simulations have been widely used to investigate the hemodynamics and mechanics of the native and prosthetic aortic valve, resolving complex flow structures, leaflet deformation, and wall shear stress (WSS).^10–17^ These studies have clarified how valve geometry, stiffness, and flow conditions affect opening kinematics and stress distribution.^18–20^ Independently, systems biology (SB) models have been developed to describe the biochemical networks and signaling pathways that regulate inflammation, fibrosis, and calcification.^21–24^ Transforming growth factor-beta (TGF-*β*) and small mothers against decapentaplegic (SMAD) signaling are known to drive fibroblast activation and extracellular matrix production,^25–27^ while endothelial NO and cyclic guanosine monophosphate (cGMP) signaling exert protective, anti-fibrotic effects by suppressing SMAD3 activity.^28, 29^

Although these mechanical and biochemical models have each contributed important insights into aortic valve calcification, they operate largely in isolation. Without coupling, they cannot capture the mechanochemical feedbacks by which mechanical dysfunction initiates molecular remodeling or how progressive fibrosis and calcification alter valve mechanics. Establishing this link is challenging because the underlying processes occur on vastly different spatial and temporal scales: valve motion and flow evolve over milliseconds, whereas biochemical signaling and tissue remodeling unfold over years. These challenges motivate the development of tractable, modular frameworks that can connect well-validated FSI solvers with mechanistic SB models to quantify mechanochemical feedback in CAVD. In this study, we present a coupled multi-physics computational framework that integrates three-dimensional FSI simulations of aortic valve dynamics with a mechanistic SB model of calcification signaling (Fig. 1). The FSI module resolves pulsatile blood flow and leaflet deformation using a sharp-interface immersed-boundary method,^16, 17, 30^ yielding spatial and temporal distributions of WSS and tissue strain throughout the cardiac cycle. These mechanical quantities serve as inputs to the SB module, which we have developed in this work by combining three major processes: (i) activation of TGF-*β* and the inflammation pathway,^31, 32^ (ii) SMAD-mediated signaling that drives fibroblastic differentiation and fibrosis,^33–35^ and (iii) NO synthesis by valvular endothelial cells (VECs) and its inhibitory effect through the NO/cGMP/protein kinase G (PKG) pathway.^6, 36, 37^ Simulation results of the coupled aortic valve calcification model reveal a self-reinforcing mechanochemical feedback: fibrosis-induced stiffening of the valve reduces WSS, decreases endothelial NO synthesis, and enhances TGF-*β* activation, thereby accelerating calcification. These findings highlight how mechanical dysfunction can amplify molecular remodeling, offering a mechanistic explanation for the rapid progression of fibrotic thickening and calcification observed in patients with CAVD.^27, 38^

**Figure 1:**
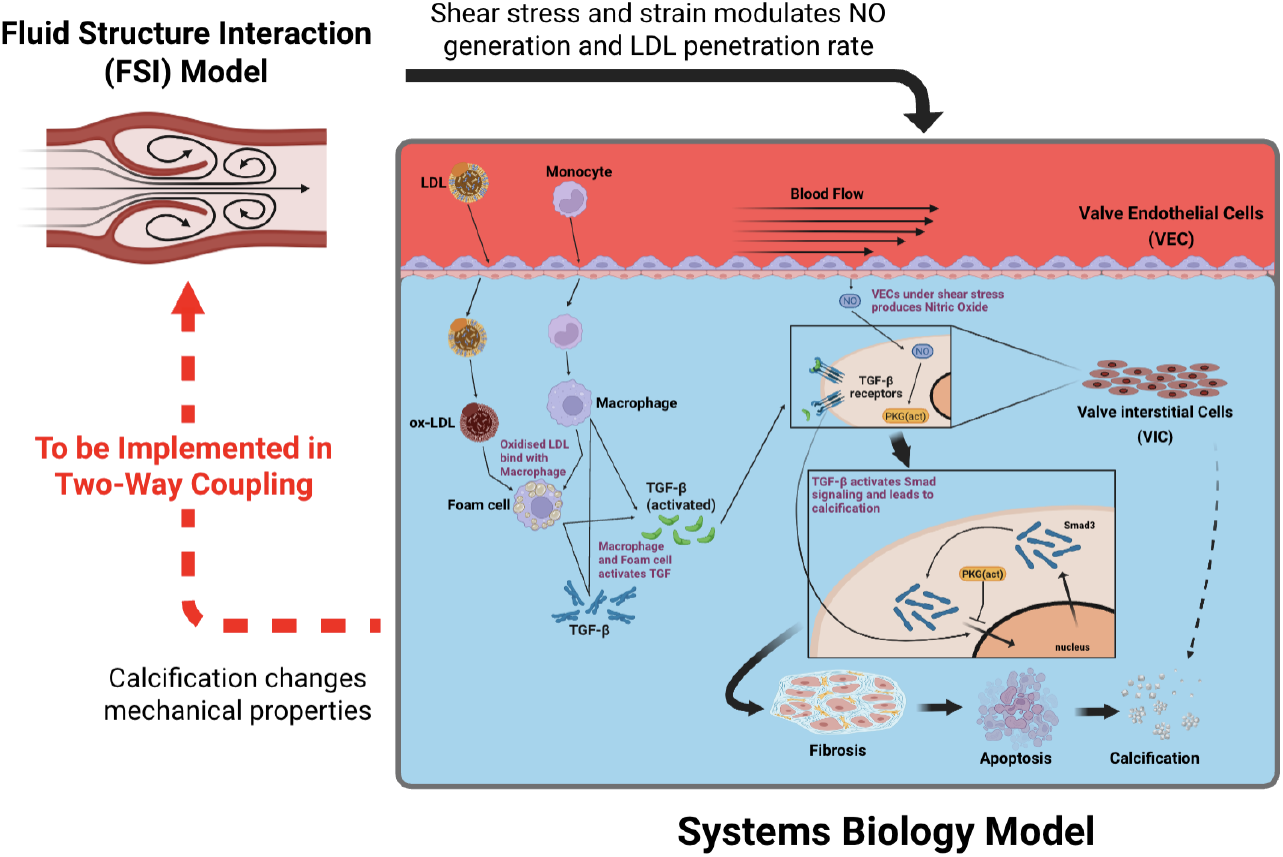
Schematic of the coupled FSI–SB framework to model mechanobiological drivers of CAVD progression. The 3D fluid structure interaction (FSI) model (left) computes flow-induced shear stress and tissue strain on the aortic valve leaflets, which serve as inputs to the systems biology (SB) model (right). These mechanical signals regulate endothelial NO production, LDL penetration, inflammatory activation, and TGF-β–dependent SMAD signaling within valve endothelial cells (VECs) and valve interstitial cells (VICs). The SB model integrates these pathways to simulate downstream fibrotic, apoptotic, and calcific processes. In the current one-way coupling implementation, information flows from the FSI model to the SB model. Future work will implement a two-way coupling scheme in which information about how changes in the tissue material properties affect blood flow through the valve (dashed red arrow on the left). NO: nitric oxide; LDL: low density lipoprotein; TGF-β: transforming growth factor β; SMAD: small mothers against decapentaplegic. Created in BioRender (biorender.com).

The multi-scale framework proposed in this work provides a foundation for exploring the coupled physical and biochemical processes that govern aortic valve remodeling and for developing predictive modeling approaches to cardiovascular disease. By linking physics-based hemodynamics with systems-level biochemistry, the framework enables dynamic simulation of how cellular signaling pathways respond to evolving mechanical environments over time. Although the present implementation employs an idealized valve geometry and the two-way coupling between the FSI and the SB module is yet to be fully implemented, these controlled simplifications allow for a computationally tractable, proof-of-principle demonstration of how hemodynamic inputs can drive biochemical remodeling *in silico*. The following sections describe the computational methodologies and model formulations used, present simulation results, and discuss the limitations, implications, and future extensions of this integrated computational approach.

## 2 Methods and Models

### 2.1 Computational FSI model of blood flow through the aortic valve

The 3D FSI model, adopted from previous work,^30^ has been applied to computational modeling of the aortic valve extensively.^16, 17^ The approach incorporates a direct forcing, immersed-boundary solver for the flow, a finite element method solver for the solid mechanics, and strong coupling between the solvers. The two solvers are parallelized through a message passing interface, so they run in parallel. At each time step, the solvers exchange data and repeat their own computation until overall convergence is achieved. Specifically, the flow solver sends the boundary load, including the pressure and shear stresses, to the solid solver, and the solid solver sends the boundary displacement and velocity of the solid body to the flow solver. Additional details of the model implementation can be found in Refs. 16 and 17.

This FSI framework is versatile and can handle complex and arbitrary geometries of solid bodies. It has previously been validated against several benchmark problems, including moving boundaries and deformable-structures with two-way interactions.^30, 39^ In the current study, we consider a simplified geometry of the aorta^17^ (Fig. 2a), where the tricuspid valve is placed within the aortic sinus with a tri-lobed dilation. The leaflets are assumed to be symmetric and have a uniform thickness, *h*, which is chosen here to be 0.3, 0.5, or 0.75 mm. The aorta is modeled as a straight and rigid tube with length *L* = 19 cm and diameter *D* = 2.24 cm. At the inlet of the tube, a transient pressure load is applied to drive blood flow (Fig. 2b). The outlet pressure is kept at 0 kPa for all simulations. This setup provides physiologically consistent results, such as the opening area, volume flow rate, and opening and closing dynamics of the valve.^17^

**Figure 2:**
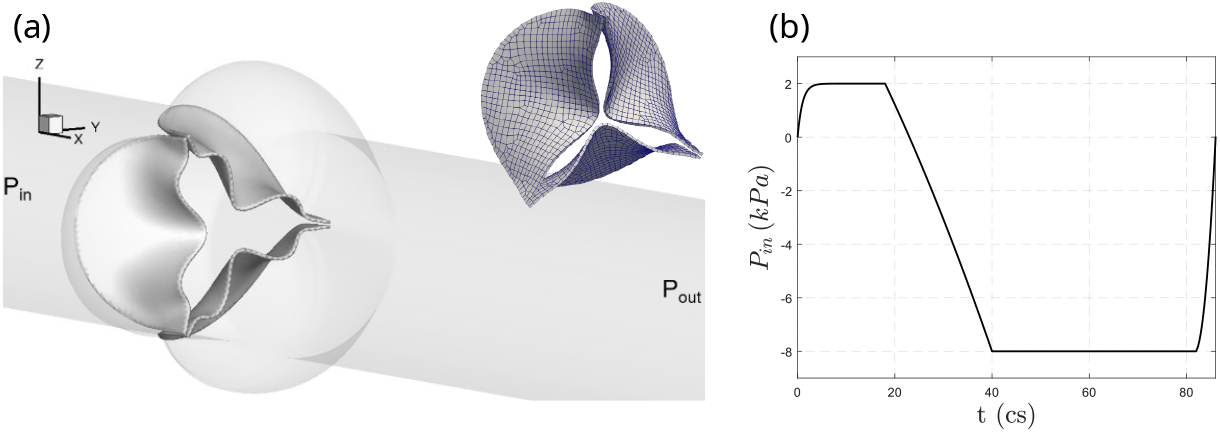
Idealized aortic valve geometry and loading conditions. (a) Geometry and finite element discretization of a tri-leaflet aortic valve attached to a cylindrical tube with three outward-shaped lobes representing the aortic root. Shown are both the anatomical configuration of the valve within the lobed tube (left) and the corresponding FEM mesh used for solid mechanics in the FSI simulations (right). (b) Prescribed transvalvular pressure waveform applied across the valve during a single cardiac cycle, with positive pressure driving systolic opening and negative pressure driving diastolic closure.

Blood is assumed to be incompressible, governed by the viscous Navier-Stokes equations,

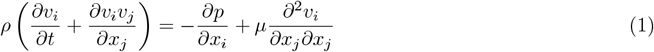

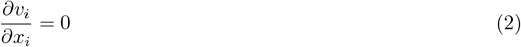

where *ρ* is blood density, *v*_*i*_ is fluid velocity, *p* is pressure, and *µ* is dynamic viscosity. Here, we set *ρ* = 1 g/cm^3^ and *µ* = 0.005 Pa·s. The leaflet tissue of the aortic valve is known to be both anisotropic and inhomogenous due to the alignment of the collagen and elastin fiber networks. For the purposes of this study, the hyperelastic Saint Venant-Kirchoff model^40^ is used to represent the behavior of the leaflets. Previous works^17^ have verified that under this model, simulated valves exhibit physiologically accurate valve opening/closing kinematics. The dynamics of leaflet deformation are governed by

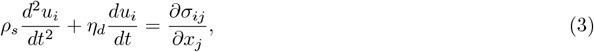

where *u*_*i*_ is the deformation, *ρ*_*s*_ is the leaflet density, *η*_*d*_ is the damping coefficient, and *σ*_*ij*_ is the Cauchy stress tensor. Here, we set *ρ*_*s*_ = 1 g/cm^3^ and *η*_*d*_ is chosen such that *hη*_*d*_ = 1 g/(cm^2^· cs). Furthermore, to calculate *σ*_*ij*_, the Young’s modulus of the tissue *E* = 2000 kPa and Poisson’s ratio *ν* = 0.4.^15, 19, 41^ A contact algorithm based on the penalty approach^16, 17^ is also applied at the leaflet surface to prevent leaflets from penetrating each other during the diastolic phase.

For spatial discretization, the aortic tube is divided into 20,735 triangle elements and 10,590 nodes, and the leaflets are divided into 1,617 20-node hexahedron elements (539 elements per leaflet; Fig. 2a). The flow domain is a 10.8 × 4.4 × 4.4 cm rectangular box, discretized by a 400 × 130 × 130 nonuniform Cartesian grid. No-slip and no-penetration boundary conditions are applied on the surface of the leaflets and the aortic wall. Simulations are run for the duration of one cardiac cycle, *T* = 0.8 s, with a flow solver timestep Δ*t*_flow_ = 4×10^−3^ centi-seconds (cs) and solid solver timestep Δ*t*_solid_ = 5 × 10^−5^ cs. Further, each Δ*t*_solid_ consists of 80 substeps.

### 2.2 SB model of aortic calcification

To capture the biochemical processes that drive CAVD progression, we constructed an integrated SB model that combines three mechanistically distinct but tightly interconnected pathways: an inflammation module describing low-density lipoprotein (LDL) infiltration, oxidation, and immune activation (Fig A1, *left*); a TGF-*β*/SMAD signaling module governing fibroblastic differentiation and pro-calcific transcriptional activity (Fig. A2); and an endothelial NO/cGMP/PKG module mediating shear-dependent inhibition of SMAD3 (Fig. A1, *right*). These sub-models, adapted from prior work^21, 29, 33^ and reformulated within a unified framework, enable dynamic simulation of how mechanical cues provided by the FSI model influence lipid transport, cytokine signaling, transcriptional regulation, and ultimately calcium deposition. The structures and key features of each sub-model and how they are coupled into a single mechanistic model of aortic valve calcification are described in the Supporting Information.

The three sub-models of the SB model were constructed and integrated in the open-source modeling and simulation platform PySB,^42^ which is designed to facilitate and streamline the construction and analysis of complex biological models. Biochemical and cellular processes are defined as reaction “rules”,^43, 44^ which can be used to generate the coupled set of ordinary differential equations (ODEs) governing the system dynamics (Eqs. S1-S46). The ODEs can then be solved numerically using integrators implemented in SciPy,^45^ which are callable from within PySB. This modular architecture is especially advantageous for our purposes, since each pathway, based on different models from the literature, uses a different unit system. By encoding each sub-model as a separate PySB module, unit consistency can be achieved programmatically, allowing for easy simulation of the full integrated SB model. The PySB model code and associated Python simulation and analysis scripts can be shared upon request.

All molecular and cellular species and biochemical reactions for the three sub-models are provided in Tables S1–S4. Initial species concentrations are provided in Table S5 and model parameter values are given in Tables S6–S8 and Fig. S1. For the inflammation pathway model (Fig. A1, *left*), concentration and rate constant units were converted from cellular concentration to volume concentration to be consistent with the units in the SMAD signaling (Fig. A2) and NO regulation (Fig. A1, *right*) models. Valvular tissue properties used for this purpose are provided in Table 1. Since the parameter values used in this study are drawn or estimated from literature sources and carry significant uncertainty, we performed a sensitivity analysis to ensure the results presented below are not significantly affected by the specific values chosen (Section 3.7).

**Table 1:** Tissue properties used to convert units of rate constants and initial conditions in the SB model. Note that the foam cell formation ratio, α, is the ratio of the rates for LDL oxidation and foam cell formation. This factor is used to convert units in the inflammation pathway to be consistent with those for the other two modules.

| Parameter | Value | Definition | Reference |
| --- | --- | --- | --- |
| $S$ | 12 cm<sup>2</sup> | Total surface area of leaflets | Current FSI model setup |
| $S_{\text{vic}}$ | 6 cm<sup>2</sup> | Total area of VICs | Current FSI model setup |
| $V_{\text{vec}}$ | $2 \times 10^{-12}$ L | Volume of a VEC | Khang et al.<sup>46</sup> |
| $\rho_{\text{vec}}$ | 10 000 cells/cm<sup>2</sup> | VEC cell surface density | Ground et al.<sup>47</sup> |
| $\rho_{\text{vic}}$ | 50 000 cells/cm<sup>2</sup> | VIC cell tissue density | Aikawa et al.,<sup>48</sup> Lis et al.<sup>49</sup> |
| $\alpha$ | $5.7 \times 10^{-6}$ cells/g | Foam cell formation ratio | Arzani et al.<sup>21</sup> |

## 3 Results

### 3.1 A multiscale framework for coupling FSI and SB models

We adopt a one-way coupling approach for the FSI-SB framework (Fig. 1). Our strategy is to first run an FSI simulation for one cardiac cycle, *T*, with assumed valve material properties to obtain spatial-temporal averaged WSS and temporally averaged maximum tissue strain. The WSS and strain information are then passed to the SB model as input to simulate the biochemical processes driving calcification. The use of only one cardiac cycle is based on the quasi-steady assumption, in which one round of FSI simulation represents a typical cardiac cycle within the time period during which the state of calcification is slow and assumed to be constant. This assumption is necessary because there is strong disparity in timescale between cycle-resolved valve mechanics (seconds) and calcification-driven remodeling (years)^21^ and it is impractical to run the expensive FSI simulations for many cycles. In the current one-way coupling framework, only the first FSI cycle is needed. To confirm that small variations in the initial WSS and strain do not significantly impact the calcification dynamics, we performed a sensitivity analysis for the initial WSS and strain values in the SB model (Section 3.7 and Figs. S2 and S3).

Both the WSS and tissue strain are calculated at each mesh node on a finite-element mesh of the valve. We use the spatial-temporal averaged WSS (SA-WSS), 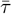, which is averaged over each side, *S*, of the leaflets over the interval *T*,

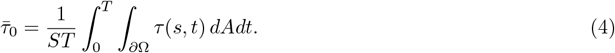

Similarly, the temporally averaged maximum tissue strain is defined as

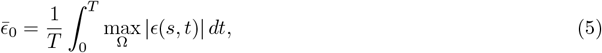

where Ω is the tissue domain and the maximum value is taken over Ω at any time point. Since we only consider one-way coupling, and thus the effects of calcification on tissue stiffness are not explicitly simulated with the FSI solver, we assume the following phenomenological relationships to update WSS and tissue strain based on calcification level, i.e.,

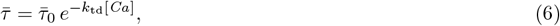

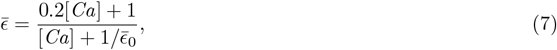

where [*Ca*] is calcification given in Agatston score.^50^ These relationships are more generalized compared to previous implementations.^21^ The updated WSS and strain feed back to calcification through Eqs. (8)–(12) (see below). In this feedback model, the SA-WSS is assumed to decrease exponentially with increasing calcification. When [*Ca*] = 0, the shear stress is simply its initial value obtained from the FSI simulation, i.e., 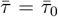. As calcium nodules grow, the resulting changes in the valve’s material properties significantly impair its ability to open, which generally reduces tissue strain. According to Eq. (7), 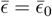 in this case. As calcification progresses and becomes severe, 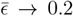, which is representative of the typical strain observed in calcified valves.^18, 51^ Although strain can exceed 0.2 in extreme cases, here we assume that calcification does not further affect strain beyond this threshold, an assumption consistent with prior modeling studies.^20, 21^ Note that these prior studies also included stress and strain updates. Here, we have generalized the approach to handle multiple cases as input variables. Additionally, *k*_td_ in Eq. (6) is used to adjust the feedback relations, and is set here to 0.006 Agatston^−1^.

In the SB model, the WSS and strain information, 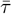 and 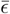, are utilized as follows:

1. The base rate constants for LDL and monocyte penetration in the absence of WSS are 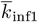 and 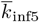, respectively. The ODEs derived from the inflammation pathway are

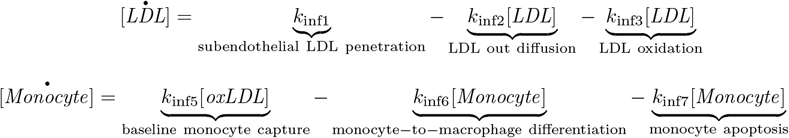

In the presence of WSS, 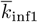 and 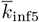 are augmented by multiplying a stress-dependent factor to reflect decreases in response,^21^ i.e.,

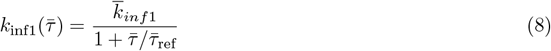

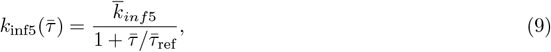

where the reference WSS, 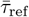, is set to 2 Pa to represent the healthy valve case. These relationships are implemented because high WSS has been shown to reinforce endothelial junctions, leading to reduced transendothelial transport of macromolecules such as LDL, and decreased adhesion of immune cells, like monocytes. This inverse relationship between shear stress and vascular permeability has been demonstrated in animal models.^52, 53^
2. The production rate of NO is WSS-dependent, giving

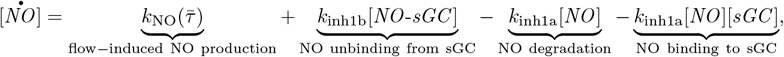

where *k*_NO_ 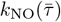 is based on data from Sriram et al.^36^ (see Fig. S1).
3. Mechanical strains in the valvular tissue amplify calcification, in part by upregulating SMAD signaling. Increased strain promotes the aggregation of valvular interstitial cells (VICs), which limits their ability to migrate and redistribute across the tissue. This localized crowding not only intensifies paracrine TGF-*β* signaling, the activator of SMAD2/3, but also predisposes VICs to apoptosis or necroptosis within these dense clusters. The resulting apoptotic bodies and matrix vesicles serve as nucleation sites for calcific nodule formation.^8^ Through this sequence, strain acts as both a mechanical and biochemical amplifier of calcification. Following a similar approach as Arzani et al.,^21^ we account for the effects of strain by assuming it directly impacts the rate constants for calcium production by SMAD2/3, *k*_sma26_ and *k*_sma27_, respectively (see Eqs. A6 and A7), according to

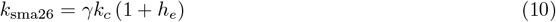

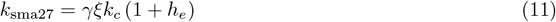

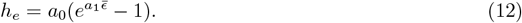

Here, *γ* is a unit conversion factor from calcium nodules/well to Agatston score (data associated with these parameters from Ref. 8 are in units of nodules/well), *k*_*c*_ is the baseline calcification rate constant [in nodules/well/(M·s)], and *a*_0_ = 4.435 × 10^4^ and *a*_1_ = 6.404 are unitless strain magnification coefficients.^21^ In the absence of strain 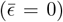, *h*_*e*_ = 0 and there is no strain magnification, giving *k*_sma26_ = *γk*_*c*_ and *k*_sma27_ = *γξk*_*c*_. Here, *ξ* is a scaling factor between 0 and 1 that represents the relative percentage of calcification by SMAD2 vs. SMAD3. Since SMAD3 is significantly more implicative of calcification,^54, 55^ we assume *ξ* = 0.15.

### 3.2 The coupled FSI–SB model predicts earlier high-risk calcification for low-average-shear, thickened aortic valves

To demonstrate the application of the coupled FSI–SB framework, we simulated long-term aortic valve calcification for three representative leaflet thicknesses: 0.3, 0.5, and 0.75 mm. For each case, SA-WSS and temporally averaged strain obtained from the FSI simulations were used to drive the biochemical dynamics governing inflammation, SMAD activation, and calcium deposition (Fig. 3). The model predicts a clear dependence of calcification rate on leaflet thickness. The thinnest leaflet (0.3 mm) shows the slowest increase in calcium accumulation, reaching an Agatston score of 400, the clinically defined high-risk threshold, at approximately 20.8 years. The 0.5-mm leaflet reaches the same threshold earlier, at roughly 18.8 years. These values fall within the range of moderate progression rates reported in clinical studies, where calcification increases steadily but remains well below the rapid progression observed in the most severely affected patients.^56^ The thickest leaflet (0.75 mm), representing a pathological case, exhibits the fastest progression, exceeding the 0.3-mm trajectory by nearly five years and reaching high-risk calcification earliest. These results illustrate how relatively small changes in leaflet thickness can lead to large differences in long-term calcification trajectories.

**Figure 3.**
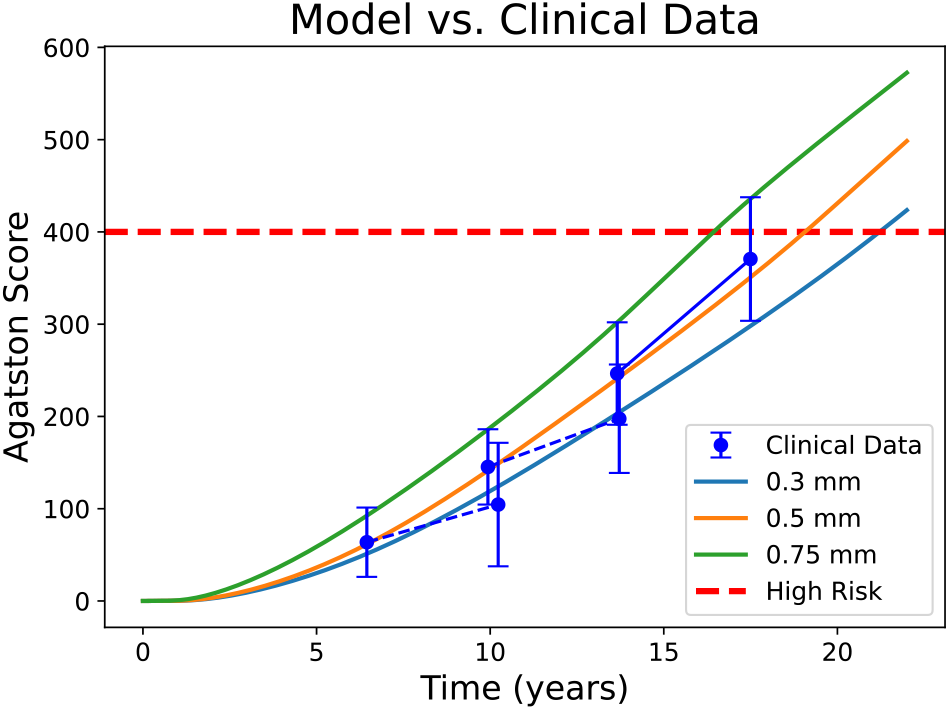
Long-term calcification trajectories predicted by the coupled FSI–SB model compared with clinical data. Model predictions for leaflet thicknesses of 0.3, 0.5, and 0.75 mm are shown over 23 years. Thinner valves accumulate calcium more slowly, while thicker valves reach the high-risk Agatston threshold years earlier. Clinical measurements from a cohort of 70 patients^56^ (solid blue line with markers) and two subgroups (lowest and mid terciles; dashed blue lines with markers) are included for comparison. The slopes of the clinical data illustrate moderate progression rates that align with the 0.3-mm and 0.5-mm model trajectories.

### 3.3 FSI simulations reveal an inverse relationship between leaflet thickness and average WSS and strain

The FSI simulations reveal systematic differences in valve kinematics, flow rate, shear stress, and tissue deformation across the three leaflet thicknesses considered (0.3, 0.5, 0.75 mm). Flow visualizations show that the 0.3-mm leaflet generates a strong, coherent systolic jet with substantial downstream vortical structures, while the 0.5-mm leaflet produces a weaker jet, and the 0.75-mm leaflet exhibits markedly reduced flow coherence due to its restricted opening (Fig. 4a). These differences are reflected quantitatively in the geometric orifice area (GOA): the 0.3-mm leaflet reaches a maximum GOA of 2.05 cm^2^, the 0.5-mm leaflet opens to 1.91 cm^2^, and the 0.75-mm leaflet achieves only 0.80 cm^2^ (Fig. 4b). Flow rate curves follow the same ordering, with peak systolic flow highest for the 0.3-mm leaflet and lowest for the 0.75-mm leaflet (Fig. 4c). These findings align with previously reported hemodynamic consequences of leaflet thickening and stiffening.^1, 2^

**Figure 4:**
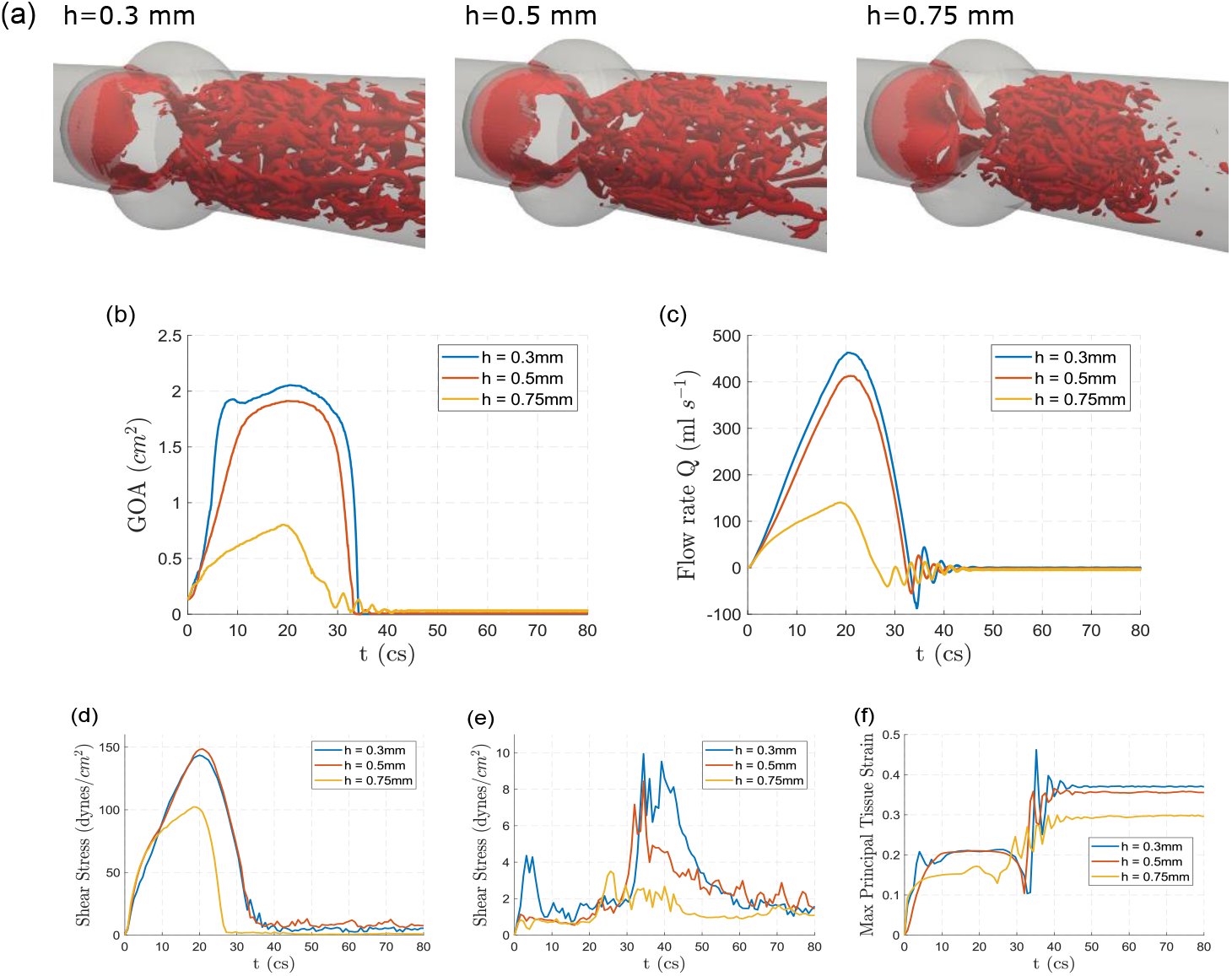
Leaflet thickening diminishes valve opening, weakens systolic flow, and reduces shear and strain. (a) Instantaneous systolic flow fields (t = 20 cs) for leaflet thicknesses of 0.3, 0.5, and 0.75 mm, visualized using the Q2-criterion, showing progressive weakening and loss of coherence in the systolic jet. (b) Geometric orifice area (GOA), where thicker leaflets not only achieve smaller peak opening but also close earlier in systole, indicating reduced mobility. (c) Flow rate curves showing diminished peak systolic flow and earlier cessation of forward flow for thicker leaflets. (d,e) SA-WSS on the ventricular (d) and aortic (e) surfaces. Ventricular-side WSS exhibits smooth temporal profiles across all three thicknesses, reflecting the predominantly unidirectional shear environment on this surface. In contrast, aortic-side WSS displays pronounced fluctuations due to its intrinsically low-magnitude, multidirectional shear environment, yet still shows a general reduction in shear levels as thickness increases. (f) Maximum principal tissue strain, showing the greatest deformation for the 0.3-mm leaflet and minimal strain for the 0.75-mm leaflet.

Differences in WSS also track consistently with leaflet thickness. Figures 4d–e show that average WSS on both the aortic and ventricular sides decreases in magnitude as thickness increases. Although wall shear stress levels are substantially higher on the ventricular side (approximately an order of magnitude greater than on the aortic side), the same thickness-dependent trend is observed on both surfaces. Spatial WSS distributions on the ventricular side further illustrate these trends: although the 0.5-mm leaflet exhibits slightly higher local shear than the 0.3-mm leaflet, reflecting its reduced bending near the attachment edge and closer alignment with the incoming jet, this difference is minimal, and the two cases remain nearly indistinguishable after spatial averaging. In contrast, the 0.75-mm leaflet shows a markedly different pattern, with high shear concentrated primarily along the free edge and much lower shear elsewhere, producing a substantially reduced average WSS due to its restricted opening and diminished flow acceleration. (Fig. 5a). Numerical values of peak and mean WSS for each case are summarized in Table 2.

**Table 2:** Key outputs from the FSI simulations. h: leaflet thickness, E_B_: bending rigidity, 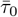: SA-WSS, 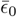: maximum strain, V_valve_: valve tissue volume, GOA_max_: maximum GOA, V_stroke_: stroke volume.

| $h$ (mm) | $E_B$ (Pa·m) | $\bar{\tau}_0$ (Pa) | $\bar{\epsilon}_0$ | $V_{valve}$ (L) | $GOA_{max}$ (cm<sup>2</sup>) | $V_{stroke}$ (mL) |
| --- | --- | --- | --- | --- | --- | --- |
| 0.3 | $5.36 \times 10^{-6}$ | 2.23 | 0.270 | $1.876 \times 10^{-4}$ | 2.05 | 88.18 |
| 0.5 | $2.48 \times 10^{-5}$ | 2.36 | 0.262 | $3.127 \times 10^{-4}$ | 1.91 | 74.81 |
| 0.75 | $8.37 \times 10^{-5}$ | 1.33 | 0.236 | $4.690 \times 10^{-4}$ | 0.80 | 19.81 |

**Figure 5:**
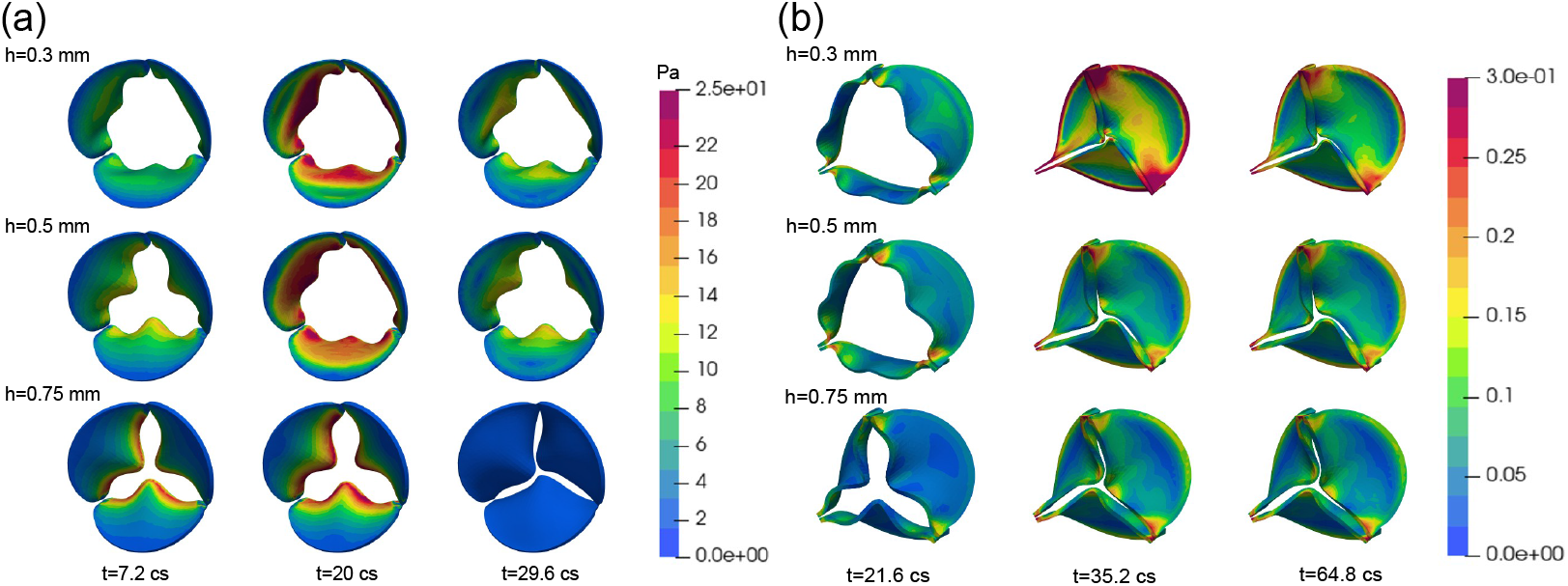
Contours of ventricular wall shear stress and aortic-side tissue strain across leaflet thicknesses. (a) Ventricular-side contours of WSS shown on the deforming valve at key systolic time points for leaflet thicknesses of 0.3, 0.5, and 0.75 mm. Thinner leaflets generate larger openings and broad regions of elevated WSS across the ventricular surface at peak systole, whereas the restricted 0.75-mm leaflet produces much lower WSS overall, with the remaining high-shear regions concentrated primarily along the leaflet-free edge due to its limited opening and reduced flow acceleration. (b) Aortic-side contours of maximum principal strain at three time points sampled throughout the cardiac cycle. Strain levels are highest during diastole, when the leaflets close and experience significant bending near the commissures and free edges. Thinner leaflets display substantially higher diastolic strain, while the thickest leaflet exhibits markedly reduced deformation across all phases.

Leaflet deformation follows the same monotonic ordering. The maximum principal strain curves in Fig. 4f and the spatial distributions in Fig. 5b show that the 0.3-mm leaflet undergoes the greatest deformation throughout systole, with high-strain regions covering substantial portions of the leaflet surface. The 0.5-mm leaflet displays intermediate deformation, whereas the 0.75-mm leaflet exhibits minimal strain. These trends are also reflected in the strain values reported in Table 2. Together, these results (Figs. 4, 5 and Table 2) demonstrate that increasing leaflet thickness consistently reduces leaflet opening, peak flow rate, WSS, and tissue strain. This establishes a clear inverse relationship between leaflet thickness and the mechanical environment that drives downstream biochemical signaling.

### 3.4 Reduced shear in thicker aortic valves decreases NO production, cGMP synthesis, and PKG activation

The long-term outputs of the NO inhibition module reflect the differences in mechanical loading established by the FSI simulations. Because endothelial NO production is strongly dependent on shear stress, the reduced WSS levels associated with thicker valves (Section 3.3) translate directly into lower concentrations of NO throughout the simulated 23-year period (Fig. 6a). The 0.3- and 0.5-mm cases exhibit similar NO trajectories during the early years of the simulation, while the 0.75-mm leaflet shows a noticeably attenuated response beginning at early time points and progressively diverging over time.

**Figure 6:**
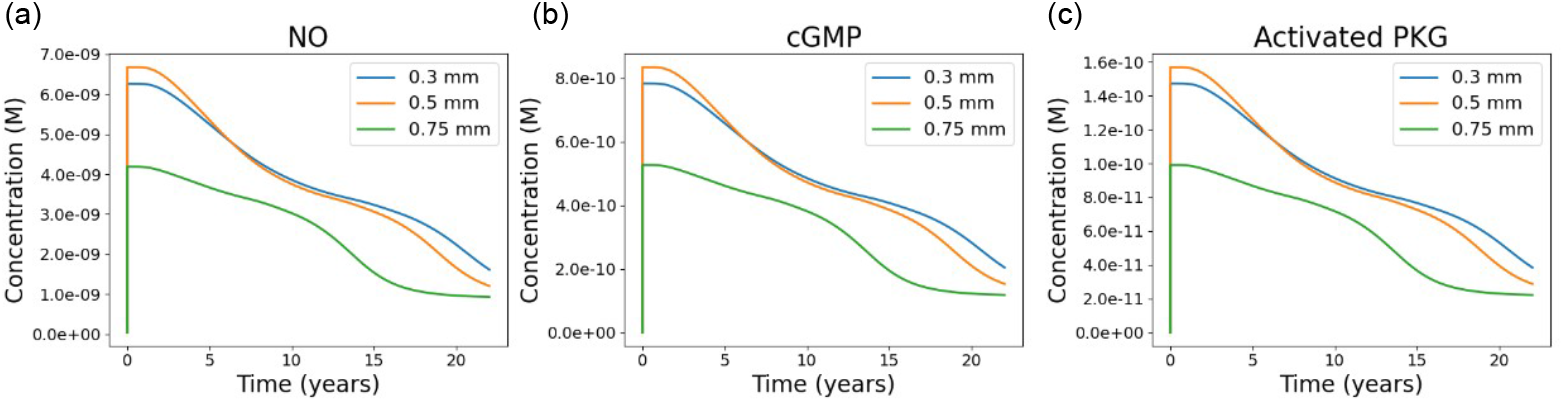
Long-term biochemical trajectories of the NO/cGMP/PKG pathway for valves of differing thickness. (a) NO concentration over 23 years for leaflet thicknesses of 0.3, 0.5, and 0.75 mm, where the thicker leaflets experience overall lower WSS, resulting in reduced NO production. (b) cGMP concentration over time, where the reduced NO levels in the thicker valves lead to lower cGMP generation. (c) Activated PKG concentrations, which depend on cGMP. Lower cGMP in the thicker valves causes PKG to activate in lower quantities as well. All three trajectories exhibit similar temporal shapes due to the tightly coupled reaction kinetics within the NO signaling module.

These reductions in NO propagate immediately to downstream components of the pathway. The cGMP concentrations produced by soluble guanylate cyclase decrease with leaflet thickness, with the thickest leaflet generating the lowest cGMP levels (Fig. 6b). Activated PKG (PKG_act_) exhibits the same trend: the 0.3- and 0.5-mm valves maintain higher PKG_act_ levels for most of the simulated duration, whereas the 0.75-mm case shows reduced activation throughout (Fig. 6c). The similar shapes of the curves for NO, cGMP, and PKG_act_ reflect the fast and tightly coupled dynamics of the NO module, where shear-driven fluctuations propagate rapidly. Taken together, these results show that variations in leaflet thickness lead to sustained differences in NO bioavailability and downstream cGMP and PKG activation. The monotonic ordering of these responses across the three thicknesses follows the ordering of WSS magnitudes from the FSI model, establishing a direct mechanochemical link between reduced shear and diminished biochemical signaling within the NO pathway.

### 3.5 Lower WSS in thicker aortic valves increases LDL and monocyte penetration and elevates downstream TGF-*β* activation

The inflammation module reveals that reductions in WSS due to increased leaflet thickness (Fig. 4d,e) lead to distinct differences in lipid infiltration and inflammatory activation over long timescales. LDL concentrations rise more rapidly and reach higher levels in the 0.75-mm leaflet compared to the 0.3-mm and 0.5-mm cases, producing an early and persistent separation between the pathological and physiological thicknesses (Fig. 7a). Although LDL levels in all cases eventually converge toward similar values by the end of the 23-year simulation, the elevated LDL burden in the early years of the 0.75-mm case establishes downstream differences in the subsequent inflammatory cascade. These differences are partly reflected in the oxidized LDL (oxLDL) dynamics. While oxLDL peaks early in all simulations, the 0.75-mm leaflet exhibits a modestly higher initial peak before all cases converge to similar levels after approximately two years (Fig. 7b). Despite the transient nature of these oxLDL differences, they are sufficient to produce noticeable changes in monocyte recruitment. Monocyte concentrations show a small early spike that is highest for the thickest leaflet case (Fig. 7c), after which they return to comparable levels across all valve thicknesses.

**Figure 7:**
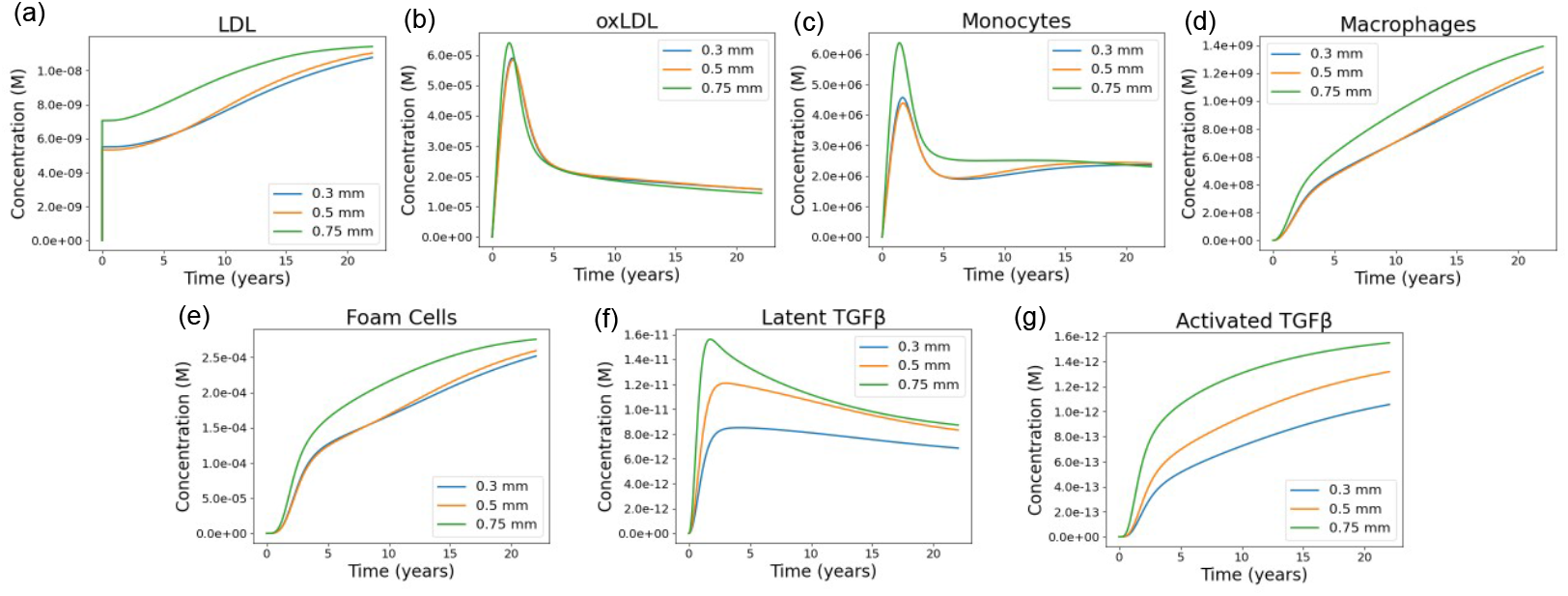
Long-term trajectories of inflammatory and TGF-β–related species for valves of differing thickness. (a, b) LDL and oxLDL concentrations over 23 years for leaflet thicknesses of 0.3, 0.5, and 0.75 mm. Lower shear in the thicker valves leads to greater LDL accumulation, producing higher oxLDL levels. For oxLDL, the three curves exhibit a short-lived difference around the 2-year mark, after which they converge, though this early divergence still propagates downstream. (c–e) Monocytes, macrophages, and foam cells, which evolve sequentially through monocyte recruitment, macrophage differentiation, and macrophage uptake of oxLDL. Monocyte levels also show a brief early separation near the 2-year peak before aligning, while macrophages and foam cells display more persistent differences, with thicker valves consistently generating larger populations due to elevated upstream oxLDL and LDL levels. (f, g) Latent and activated TGF-β concentrations, produced downstream of foam cell activity. Latent TGF-β trajectories partially converge after their early peak, but activated TGF-β maintains clear separation across leaflet thicknesses, reflecting sustained differences in upstream inflammatory signaling.

In contrast to monocytes, the macrophage and foam cell populations show large and persistent differences among the three cases. Both populations increase throughout the simulated period, but the 0.75-mm leaflet maintains substantially higher macrophage (Fig. 7d) and foam cell (Fig. 7e) levels than the 0.3-mm and 0.5-mm cases across the entire 23-year timeframe. This sustained divergence arises despite the relatively small and short-lived differences in oxLDL and monocyte levels, highlighting the sensitivity of the inflammatory module to early perturbations in lipid infiltration. The effects of these inflammatory differences propagate to the TGF-*β* subsystem. Latent TGF-*β* rises more quickly in the 0.75-mm case and reaches a higher early peak before gradually declining and converging with the 0.5-mm case (Fig. 7f). Activated TGF-*β*, however, shows a persistent separation across all cases, with the 0.75-mm leaflet maintaining the highest levels throughout the simulation, followed by 0.5-mm and then 0.3-mm (Fig. 7g). Because macrophages and foam cells contribute to TGF-*β* activation, their sustained elevation in the thickest-leaflet case likely drives the long-term divergence observed in activated TGF-*β*. These results show that even modest, short-lived differences in LDL infiltration and oxLDL exposure can amplify through the nonlinear inflammatory cascade, producing long-lasting differences in macrophage and foam cell populations and, ultimately, sustained elevation of activated TGF-*β* in thicker, low-shear valves.

### 3.6 Thicker valves exhibit increased total SMAD phosphorylation and reduced SMAD3 inhibition, leading to enhanced pro-calcific signaling

The SMAD signaling module integrates the combined biochemical inputs from the upstream pathways driven by WSS, NO availability, and TGF-*β* activation. Because total phosporylated SMAD (pSMAD2/3) formation depends on active TGF-*β* (TGF-*β*_act_) while phosphorylated SMAD3 (pSMAD3) inhibition depends on active PKG (PKG_act_), the SMAD responses reflect the trends established in the mechanical (Figs. 4 and 5), NO inhibition (Fig. 6), and inflammation (Fig. 7) results. The model captures the expected SMAD signaling behavior in response to these thickness-dependent upstream cues (Fig. 8). Nuclear pSMAD2 concentrations rise monotonically across all cases, with the 0.75-mm leaflet producing the highest pSMAD2 levels due to its elevated TGF-*β*_act_ concentrations (Fig. 8a). Cytoplasmic pSMAD2 exhibits a similar ordering (Fig. 8c). Nuclear pSMAD3 also displays an amplified response in the thickest leaflet, but with a distinct transient peak followed by a decline (Fig. 8b). This peak arises from the combined influence of higher TGF-*β*_act_ levels (Fig. 7g) and lower PKG_act_ levels (Fig. 6c). The cytoplasmic pSMAD3 profiles follow the same pattern (Fig. 8d).

**Figure 8:**
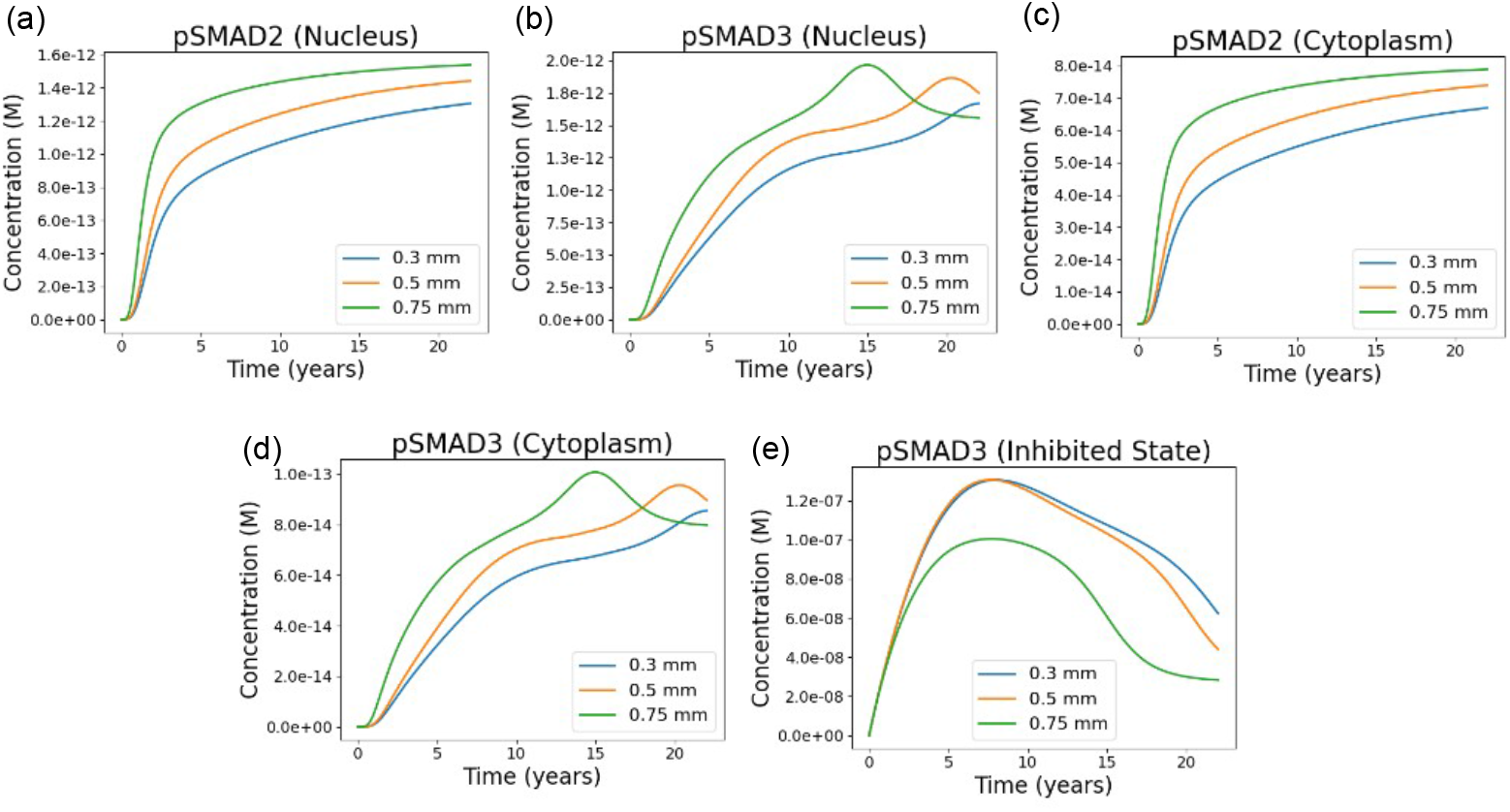
Long-term trajectories of SMAD2/3 signaling species for valves of differing thickness. (a, b) Nuclear pSMAD2 and pSMAD3 concentrations over 23 years for leaflet thicknesses of 0.3, 0.5, and 0.75 mm. Nuclear pSMAD2 increases and then approaches a steady state, with thicker valves maintaining higher levels due to their elevated TGF-β activation. In contrast, nuclear pSMAD3 exhibits a transient peak followed by a decline, reflecting the competing influences of TGF-β–driven phosphorylation and PKG-mediated inhibition. The thickest valves sustain the highest overall pSMAD3 levels due to stronger TGF-β signaling and weaker PKG activity. (c, d) Cytoplasmic pSMAD2 and pSMAD3 show the same distinction: pSMAD2 rises toward a steady-state plateau, whereas pSMAD3 displays a peak-and-decline profile driven by the same balance between TGF-β and PKG. (e) Inhibited pSMAD3, produced through PKG-dependent hyperphosphorylation, is highest in the thinner valves, reflecting their stronger NO/cGMP/PKG signaling, and lowest in the thickest valve, consistent with diminished PKG activity and sustained TGF-β exposure.

The inhibited pSMAD3 trajectories further illustrate the importance of PKG-mediated repression of SMAD3. The 0.3- and 0.5-mm cases show higher inhibited pSMAD3, reflecting their stronger NO/PKG responses (Fig. 6) and lower TGF-*β*_act_ concentrations (Fig. 7g). In contrast, the 0.75-mm leaflet exhibits markedly reduced SMAD3 inhibition (Fig. 8e), consistent with both diminished PKG_act_ and sustained TGF-*β*_act_ production. These results show that thicker aortic valve leaflets exhibit increased total SMAD2/3 phosphorylation and reduced PKG-dependent SMAD3 inhibition. The resulting elevation in transcriptionally active pSMAD3 provides a direct pro-calcific signal within the integrated signaling network, connecting the upstream mechanical and biochemical differences (Figs. 4–7) to the accelerated calcification trajectory observed for thicker valves (Fig. 3).

### 3.7 Sensitivity analysis

A total of 73 parameters were included in the sensitivity analysis following a standard approach.^57^ This includes 12 rate constants from the inflammation pathway model (Table S6), 39 rate constants from the SMAD signaling model (Table S7), 15 rate constants from the NO regulation model (Table S8), *ξ* from Eq. (11), *a*_0_ and *a*_1_ from Eq. (12), *b*_0_ and *b*_1_ from Eq. S49, and 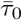 and 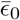, the initial values of WSS and strain obtained from the FSI simulator, in Eqs. (4) and (5), respectively. Parameter values were varied by *±*10%, while the mechanical inputs were varied by *±*15%, and the relative change in the calcification level at *t* = 23 years was recorded. More details are provided in Supplementary Material. For the FSI-based inputs 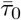 and 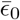, we see *<* 5% changes in calcification level in all cases (Figs. S2 and S3). For the other 71 parameters, changes in calcification level were *<* 25% in all cases, with only eight parameters producing changes *>* 10% (Figs. S4 and S5). For the purposes of this proof-of-principle demonstration of our framework, these results confirm that the general trends reported are not significantly impacted by the parameter values used.

## 4 Discussion

We present a mechanochemical framework that couples 3D FSI with systems-level biochemical signaling to model CAVD progression. By supplying FSI-derived shear stresses and tissue strains directly to an integrated SB model of the biochemical and cellular processes driving aortic calcification (Fig. 1), the coupled FSI-SB framework captures how differences in the mechanical environment translate into divergent long-term calcification trajectories. In prior mechanobiological models of CAVD,^21^ baseline shear stress and strain were informed by FSI simulations reported in the literature^58^ and by experimental measurements.^59^ Subsequent evolution of these mechanical cues was governed by phenomenological relationships tied to a single baseline mechanical state. In contrast, the present framework computes baseline shear stress and strain directly using our custom FSI simulator^16, 17, 30^ and treats these quantities as explicit inputs to the SB model. The evolution of shear stress and strain is then governed by generalized relationships (Eqs. 6 and 7) that are not calibrated to a specific reference state. This enables systematic exploration of how valve properties, such as leaflet thickness, alter signaling and calcification trajectories. Furthermore, the SB model used here extends the model developed in prior work^21^ by coupling inflammation processes to NO regulation^29, 60–62^ and TGF-*β*/SMAD signaling.^28, 33, 54, 55, 63–65^ These pathways play central roles in transducing shear- and strain-dependent signals at the endothelium, regulating VIC activation, and initiating fibroblastic differentiation. Incorporating NO and SMAD signaling enables the SB model to capture key regulatory feedbacks linking mechanical stimuli to downstream calcification processes that cannot be represented by inflammation-driven mechanisms alone.

As a proof-of-principle demonstration of the FSI-SB platform, we quantified how leaflet thickness alters flow-induced mechanical forces and biochemical signaling in aortic valves, resulting in divergent long-term calcification. Simulations show that thinner, more compliant valves maintain higher shear and stronger NO-mediated inhibition, while thicker, fibrotic valves exhibit reduced shear, enhanced TGF-*β* activation, and accelerated calcification (Fig. 3). FSI results reveal that thinner leaflets maintain larger orifice areas, stronger flow jets, higher WSS, and greater tissue strains, whereas thicker leaflets open less and experience systematically reduced mechanical loading (Figs. 4, 5, and Table 2). At the biochemical level, reduced WSS leads to lower NO levels and diminished cGMP and PKG activation over a *>* 20-year simulated period (Fig. 6), consistent with prior reports that shear-dependent NO signaling suppresses early pro-calcific activity.^6^ Lower WSS also increases LDL penetration (Fig. 7a), producing transient differences in oxidized LDL and monocyte recruitment (Fig. 7b,c). This causes persistent increases in macrophages and foam cells (Fig. 7d,e), in agreement with clinical and experimental observations that inflammatory loading accelerates CAVD progression.^5^ These immune cell populations drive higher TGF-*β* activation in thicker leaflets (Fig. 7f,g), which is then amplified by the SMAD pathway. Cytoplasmic and nuclear phosphorylated SMAD2 increase across all cases (Fig. 8a,c), while phosphorylated SMAD3 exhibits an early peak due to competing influences of TGF-*β* and PKG (Fig. 8b,d). Reduced PKG-mediated inhibition in thicker valves results in a larger pool of active phosphorylated SMAD3 (Fig. 8e), consistent with reports linking low shear and diminished NO signaling to accelerated calcification and poor clinical outcomes.^6^ These interconnected mechanobiological effects collectively explain the observed divergent calcification trajectories (Fig. 3).

Despite these encouraging results, several limitations of the present FSI-SB framework should be noted. First and foremost, the current implementation uses a *one-way coupling* in which FSI-derived WSS and strain drive biochemical dynamics, but mechanical consequences of tissue calcification do not feed back to update the tissue stiffness or FSI (Fig. 1). Instead, phenomenological adjustments are used to approximate stiffening effects in an efficient manner (Eqs. 6 and 7). A fully *two-way coupling*, physics-based implementation would mitigate this limitation by periodically feeding the predicted calcification state from the SB model back to the FSI model to update the leaflets’ mechanical properties and recompute WSS and strain (Fig. 1). This feedback can be done while retaining the phenomenological update laws between FSI updates, capitalizing on the short-term accuracy and efficiency of these relations. Implementing such an iterative, multiscale coupling strategy will reduce reliance on the update laws over multi-decade simulations by recalibrating the mechanical environment at intermittent intervals or when calcification changes exceed a prescribed threshold. Quantitative accuracy and long-term reliability of model predictions are thus expected to improve by more faithfully capturing the evolving mechanical–biochemical feedback that influences calcification progression.

Additional limitations of the present FSI–SB framework include: (i) a simplified tri-leaflet valve geometry in the FSI solver, coupled to a straight aortic channel with prescribed material properties; (ii) the use of spatially-averaged, leaflet-level representations of WSS, strain, and calcification; and (iii) our reliance in the SB model on uncertain parameters from the literature, which in many cases lack accompanying experimental measurements in the context of valve calcification. For the first issue, a simplified geometry was intentionally chosen for the proof-of-principle application presented here because it allowed us to isolate thickness-dependent mechanical effects. However, since WSS and leaflet strain depend strongly on valve geometry and material properties, future work will utilize the full suite of features supported by our FSI solver,^30^ including alternative tube and leaflet shapes and variations in material properties. To address the second issue, the integrated SB model will be formulated within a reaction-diffusion framework to incorporate spatially-resolved signaling, regional variations in mechanical stimuli, and heterogeneous VIC phenotypes. This feature will allow us to utilize the full spatially-resolved shear stress and strain fields computed by the FSI solver. Finally, to assess the impact of parameter uncertainty on the results of our study, we performed a sensitivity analysis of terminal calcification outcomes (Figs. S2–S5). Future studies will improve upon this analysis by performing formal parameter calibration and uncertainty quantification using Bayesian inference approaches,^66, 67^ enhancing quantitative accuracy and predictive confidence.

A major advantage of the present FSI-SB framework is its modular architecture, which provides significant flexibility and opportunities for future development. The SB model can be readily expanded to include additional pathways implicated in CAVD, such as bone morphogenetic protein signaling and SMAD1/5/7 regulation,^68^ matrix metalloproteinase activity,^69^ NO-dependent S-nitrosylation,^70^ and osteogenic differentiation of VICs.^71^ The platform also supports simulation of pharmacological interventions by targeting specific biochemical reaction branches, enabling *in silico* exploration of mechanosensitive therapies. To validate the extended FSI-SB framework, three experimental strategies can be pursued. First, *in vitro* flow chamber studies using VICs and VECs embedded in 3D collagen or hydrogel matrices can be used to collect molecular-level data such as NO production, TGF-*β*/BMP signaling, and expression of osteogenic markers like RUNX2 and ALP under controlled shear and strain.^6, 72^ Second, animal models, including mouse and porcine, may be adopted to enable measurements of both leaflet-level biomechanics (e.g., strain, shear) and physiological outcomes over time, including calcified nodule formation, histological remodeling, and early osteochondral gene expression.^73^ Finally, additional longitudinal clinical imaging, such as serial CT-derived Agatston scores and Doppler echocardiography, may be used to assess whether the model accurately predicts real-world calcification rates and functional valve deterioration.^74–77^ Overall, the present work establishes a foundation for mechanobiological modeling of CAVD progression, with the long-term goal of supporting personalized diagnostics, risk stratification, and therapeutic design.

## 5 Conclusion

Computational modeling is becoming an essential tool for understanding and ultimately treating CAVD. As experimental techniques increasingly reveal the molecular and mechanical complexity of CAVD, there is a growing need for integrative frameworks capable of unifying these multiscale processes into coherent, predictive computational models. The framework developed in this study represents a step toward that goal by linking FSI with mechanistic biochemical signaling to simulate long-term disease progression. By capturing how changes in aortic leaflet mechanics reshape endothelial signaling, inflammatory activation, and SMAD-mediated transcription, the model illustrates the power of mechanochemical simulations to uncover pathways through which mechanical remodeling influences cellular decision-making. Although the present work focuses on leaflet thickening, the framework is sufficiently flexible to incorporate patient-specific geometries, emerging molecular pathways, and pharmacological perturbations. With future extensions, including two-way coupling, spatially resolved VIC dynamics, and calibrated biochemical parameter sets, the framework can support predictive analyses of disease progression and provide a platform for virtual testing of therapeutic interventions. As computational cardiology continues to advance, models of this type will play an increasingly central role in identifying early mechanobiological indicators of disease, evaluating potential treatment strategies, and personalizing care for patients with aortic valve pathology. The approach developed here demonstrates how high-fidelity FSI simulations and SB models can be integrated to generate mechanistic insight and inspire new strategies for preventing or slowing CAVD.

## 6 Supporting Information

The Supporting Information (SI) accompanying this manuscript provides a comprehensive description of the model formulation and implementation. It includes a complete listing of all biochemical reactions incorporated in the signaling network and the corresponding system of ordinary differential equations derived from them. Detailed parameter documentation is also provided, including rate constants, amplification factors, and other kinetic terms, along with their physiological justification and cited literature sources. Furthermore, the SI specifies the initial conditions used for each molecular species, their biological rationale, and references to experimental data from which these values were obtained. Together, these materials ensure full transparency and reproducibility of the computational model.

## Acknowledgments

This work is funded by the National Science Foundation Graduate Research Fellowship Program (NSF-GRFP) to MQ (Award # 1937963 and 2444112) and a National Institutes of Health K22 Career Development Award (K22-CA37857) to LAH. The 3D FSI simulations were performed on the Stampede3 supercomputer at the Texas Advanced Computing Center (TACC) through the *Advanced Cyberinfrastructure Coordination Ecosystem: Services and Support (ACCESS)* program.

## Appendix Biochemical pathways in the systems biology model

### Inflammation pathway

**Figure A1:**
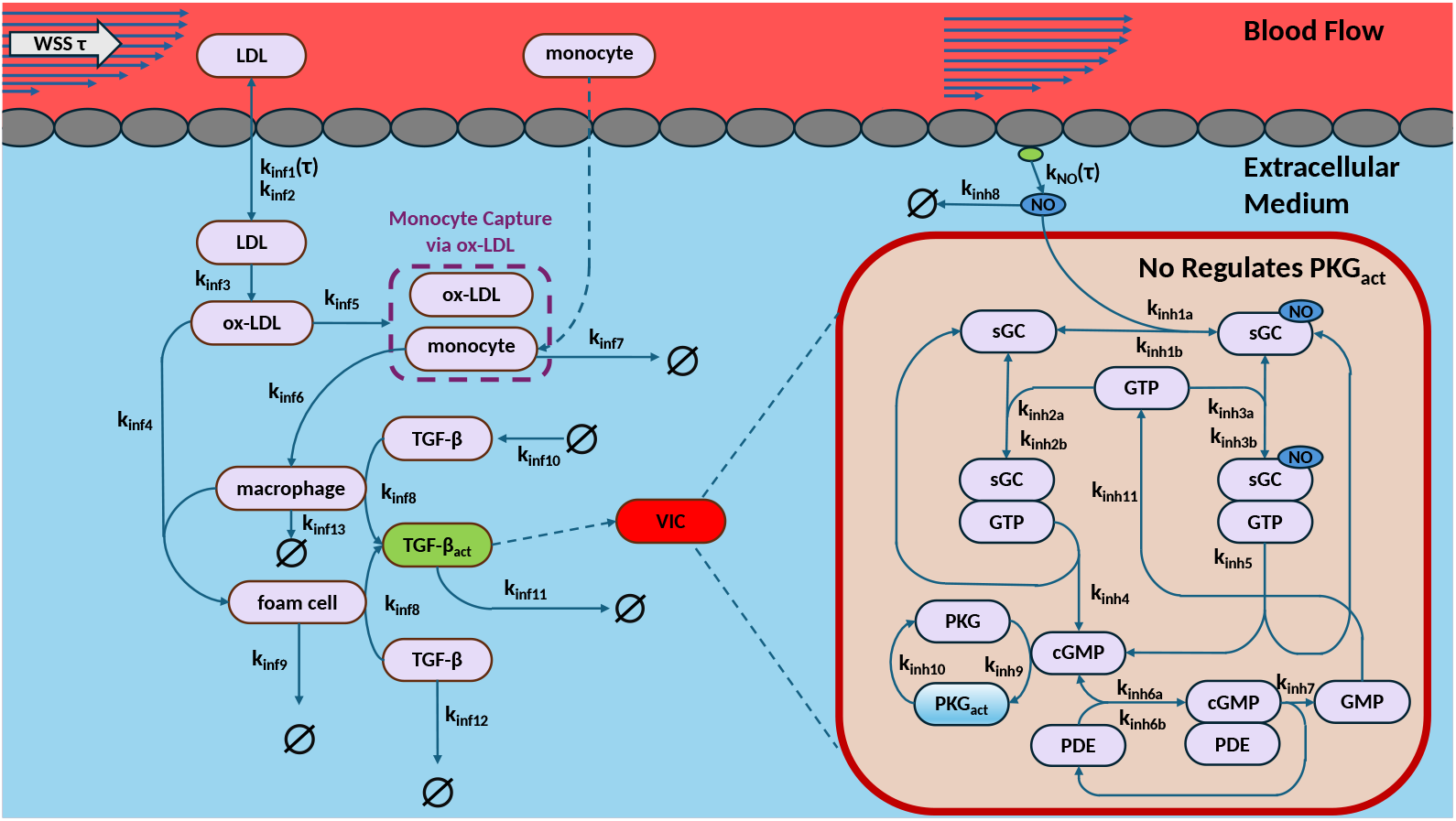
Mechanistic models of inflammation and NO regulation in aortic valve calcification. In the inflammation model (left), low WSS increases subendothelial LDL penetration and oxidation, promoting monocyte recruitment, macrophage and foam cell formation, and activation of latent TGF-β. Active TGF-β serves as the upstream input to VIC SMAD signaling. The NO pathway (right) describes WSS-dependent NO production via eNOS, subsequent sGC-mediated cGMP generation, and PKG activation, which inhibits pro-calcific SMAD3 signaling. Arrows denote biochemical reactions; dashed arrows represent transport or recruitment processes.

Since fibroblastic differentiation and subsequent apoptosis of valve interstitial cells (VICs) are a predominant contribution to the formation of calcific nodules on aortic valve leaflets,^4^ the SB inflammation model, adapted from Arzani et al.,^21^ focuses on calcification mediated through fibroblastic pathways (Fig. A1). The process begins with the subendothelial infiltration of low-density lipoproteins (LDL) into the valve tissue, where LDL undergoes oxidation. Oxidized LDL (oxLDL) promotes the recruitment of circulating monocytes from the bloodstream, which differentiate into macrophages within the valve’s microenvironment.^1^ Macrophages can further interact with oxLDL to form foam cells, which are lipid-laden cells implicated in chronic inflammation and commonly observed in atherosclerotic plaques. This process is modeled as

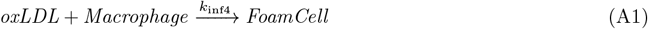

Accumulation of immune cells within the tissue, particularly macrophages and foam cells, leads to the activation of latent TGF-*β*, initially secreted by macrophages in an inactive form.^78, 79^ These activation processes are modeled as

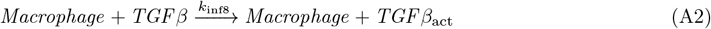

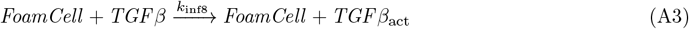

Once activated, TGF-*β* binds to receptors on the surface of VICs, initiating the process of myofibroblastic differentiation.^78^ This transition promotes ECM remodeling, VIC apoptosis, and ultimately the formation of calcific nodules. Note that the Arzani et al.^21^ inflammation model does not include SMAD signaling, with calcification directly dependent on TGF-*β*. Macrophages are also assumed to grow without bound. Here, we remedy this by including macrophage cell death, modeled as [*Macrophage*] 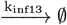

#### SMAD signaling

SMAD signaling is initiated within VICs upon binding of activated TGF-*β* to both TGF-*β* receptor types I and II (TGF*β*RI and TGF*β*RII) on the VIC surface (Fig. A2). We model TGF-*β* ligand-receptor binding by

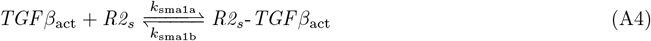

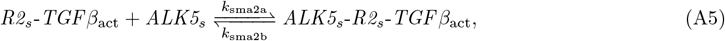

where ALK5 is a known TGF*β*RI and R2 is a generic TGF*β*RII. The resulting ALK5-R2-TGF*β*_act_ receptor complex translocates to the cytoplasm, where it phosphorylates SMAD2 and SMAD3 signaling proteins. These phosphorylated SMADs then heterodimerize with SMAD4 to form the SMAD transcription factor complex.^80, 81^ Upon nuclear translocation, this complex promotes transcription of pro-fibrotic genes, driving fibroblastic differentiation of the VICs. Continued expression of these genes, including those involved in excessive extracellular matrix (ECM) production and overexpression of *α* smooth muscle actin (*α*SMA), leads to apoptosis. This process creates nucleation sites for calcium deposition, allowing hydroxyapatite, i.e., calcification, nodules to form.^82, 83^

The SMAD signaling pathway model used in this work is adapted from Chung et al.^33^ However, to fully capture the dynamics of SMAD signaling, we have modified the Chung et al. model to explicitly include SMAD3. Even though SMAD2 and SMAD3 exhibit similar cytoplasmic kinetics,^63^ their roles diverge downstream: SMAD3 more strongly influences fibroblastic differentiation and ECM production,^54, 55^ making it the primary driver of the calcification response. Accordingly, following prior work by Arzani et al.,^21^ we model calcification (*Ca*) as a first-order process driven by the SMAD transcription factor complex, with a rate constant that varies depending on whether the complex contains SMAD2 or SMAD3, i.e.,

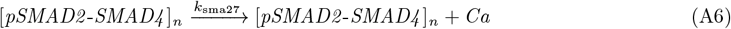

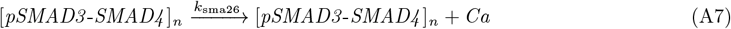

Furthermore, the SMAD signaling pathway has an internal autoregulatory mechanism, i.e., the NO/cGMP/PKG pathway.^64, 65^ Activation of PKG by this pathway leads to hyperphosphorylation of SMAD3, inhibiting heterodimerization with SMAD4 and preventing translocatation to the nucleus.^28^ Here, we model SMAD3 hyper-phosphorylation as a bimolecular process,

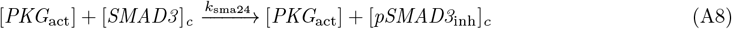

#### Endothelial NO regulation

Counteracting the pro-fibrotic SMAD pathway, the NO/cGMP/PKG signaling cascade, mediated by valvular endothelial cells (VECs), is initiated in response to high shear stress (Fig. A1). Upon valve opening, VECs lining the surface of the valve leaflets, particularly on the ventricularis (the side facing the left ventricle), experience elevated shear stress. This mechanical stimulus activates endothelial NO synthase (eNOS), leading to the production of NO,^6, 65^ which diffuses into the underlying VICs. There, it activates soluble guanylyl cyclase (sGC), catalyzing the conversion of guanosine triphosphate (GTP) to cGMP.^29^ The NO regulation model used here is adapted from Garmaroudi et al.,^29^ which includes sGC activation by NO and production of cGMP. In this work, we add two additional reactions that follow cGMP production: autoinhibitory PKG deactivation,^60^ modeled as the first-order reaction [*PKG*] 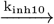 [*PKG* ], and recycling of GMP back to GTP.

Although a multifaceted process,^61, 62^ for simplicity, here we model GMP recycling as the first-order process [*GMP* ] 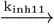 [*GTP* ].

As a secondary messenger, cGMP induces smooth muscle relaxation by activating cGMP-dependent protein kinases, such as PKG, modeled here as

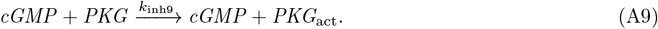

**Figure A2:**
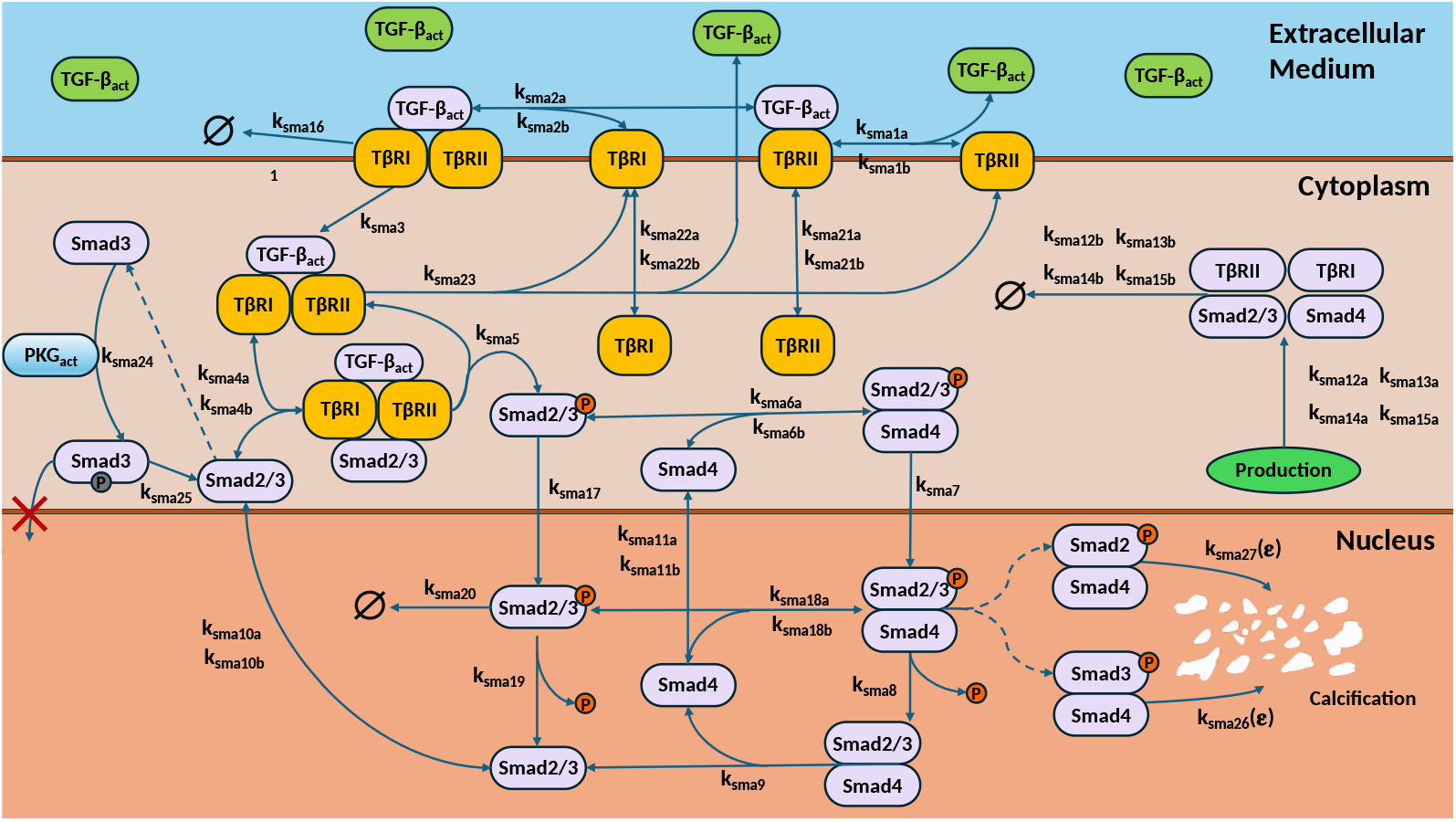
Mechanistic model of TGF-β–SMAD signaling leading to VIC calcification. Active TGF-β binds sequentially to TGFβRII and TGFβRI at the VIC surface, forming a receptor complex that phosphorylates cytoplasmic SMAD2 and SMAD3. Phosphorylated SMADs heterodimerize with SMAD4 and translocate to the nucleus, where they promote pro-calcific gene expression. SMAD2 and SMAD3 are treated with identical kinetics except for PKG-dependent hyperphosphorylation of SMAD3, which prevents its association with SMAD4. Active PKG (PKG_act_) is supplied by the NO pathway (Fig. A1), providing the biochemical link through which WSS-dependent NO production inhibits SMAD3-driven calcification. Arrows denote biochemical reactions, and dashed arrows represent molecular transport between compartments.

Activated PKG then inhibits pro-calcific signaling pathways, including those driven by SMAD3^60^ (see Eq. (A8)). Although this pathway mitigates the progression of calcific nodule formation, its cardioprotective effects are limited. Persistent ECM production by quiescent VICs, along with leaflet damage that facilitates increased LDL and monocyte infiltration, compromises the effectiveness of NO signaling. These factors contribute to progressive valve leaflet thickening, which alters the shear stress environment and further impairs NO-mediated protection.^9^

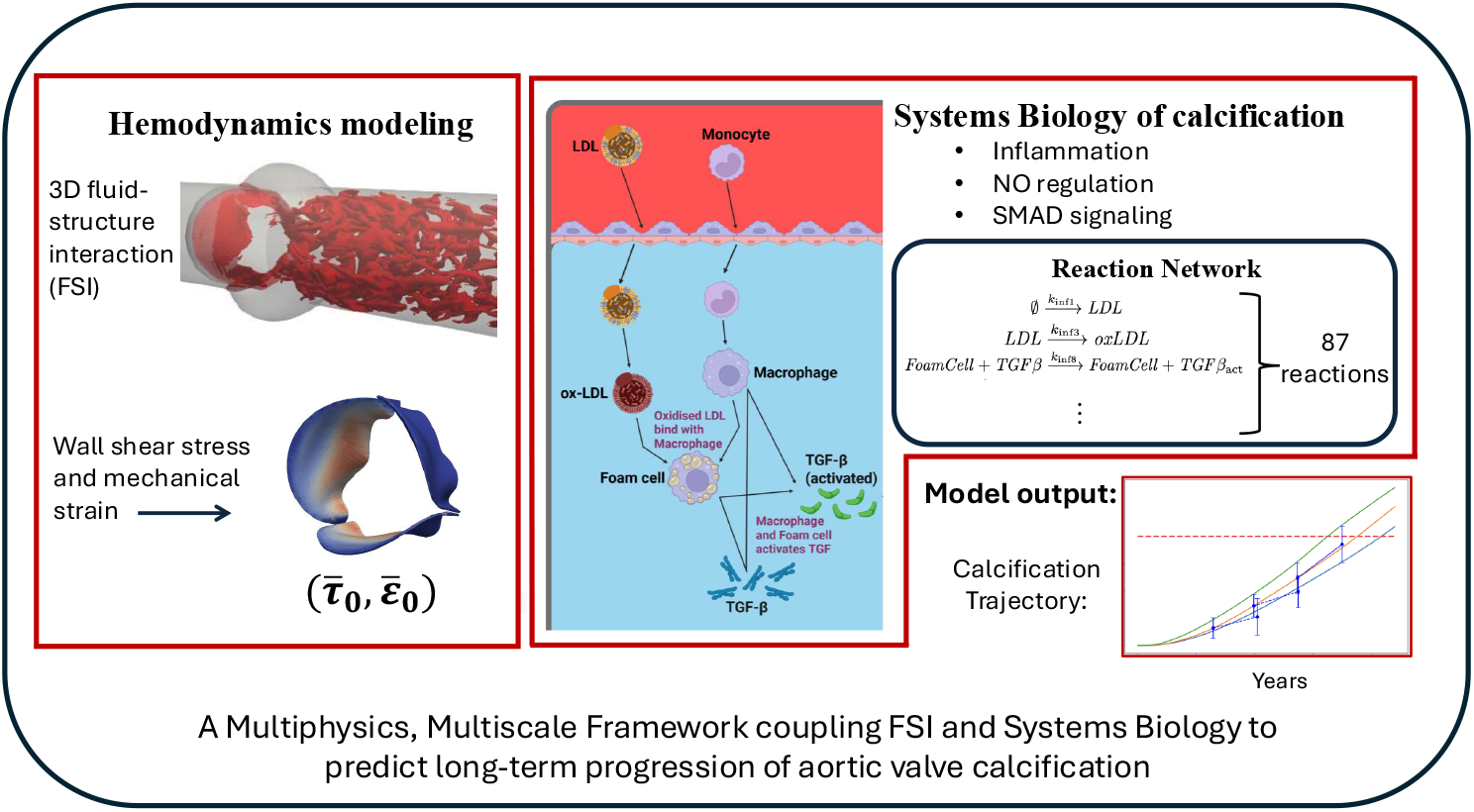

