## Supplemental Text for "Capturing Multi-Scale Dynamics of Aortic Valve Calcification With a Coupled Fluid–Structure and Systems Biology Model"

### Supporting Information

#### Species in the integrated SB aortic calcification model

**Table S1:** ***List of species***

| Species | Description |
|---|---|
| $ALK5_c$ | Cytoplasmic ALK5 (TGF$\beta$ receptor type I) |
| $ALK5_c\text{-}R2_c\text{-}TGF\beta_{\mathrm{act}}$ | Cytoplasmic ALK5–R2–TGF$\beta_{\mathrm{act}}$ receptor complex |
| $ALK5_s$ | Surface ALK5 (TGF$\beta$ receptor type I) |
| $ALK5_s\text{-}R2_s\text{-}TGF\beta_{\mathrm{act}}$ | Surface ALK5–R2–TGF$\beta_{\mathrm{act}}$ receptor complex |
| *Ca* | Intracellular calcium produced downstream of SMAD-dependent transcription |
| *FoamCell* | Foam cell (lipid-loaded macrophage) |
| *sGC* | Soluble guanylyl cyclase |
| *sGC-GTP* | sGC–GTP complex |
| *GMP* | Guanosine monophosphate produced by PDE hydrolysis of cGMP |
| *cGMP* | Cyclic GMP, second messenger produced by sGC |
| *GTP* | Guanosine triphosphate, substrate for sGC catalytic activity |
| *LDL* | Low-density lipoprotein |
| *oxLDL* | Oxidized LDL |
| *Macrophage* | Differentiated monocyte involved in inflammation and TGF$\beta$ activation |
| *Monocyte* | Circulating immune cell that differentiates into macrophages |
| *NO* | Nitric oxide |
| *NO-sGC* | NO–sGC complex |
| *NO-sGC-GTP* | NO–sGC–GTP complex |
| *PDE* | Phosphodiesterase enzyme |
| *PDE-cGMP* | PDE–cGMP complex |
| *PKG* | Inactive protein kinase G |
| $PKG_{\mathrm{act}}$ | Active PKG (activated by cGMP) |
| $R2_c$ | Cytoplasmic TGF$\beta$ receptor type II |
| $R2_s$ | Surface TGF$\beta$ receptor type II |
| $R2_s\text{-}TGF\beta_{\mathrm{act}}$ | Surface R2–active TGF$\beta$ complex |
| $SMAD2_c$ | Cytoplasmic SMAD2 |
| $pSMAD2_c$ | Cytoplasmic phosphorylated SMAD2 |
| $SMAD2_c\text{-}ALK5_c\text{-}R2_c\text{-}TGF\beta_{\mathrm{act}}$ | Cytoplasmic TGF$\beta$ receptor complex with SMAD2 and active TGF-$\beta$ ligand |
| $pSMAD2_c\text{-}SMAD4_c$ | Cytoplasmic pSMAD2–SMAD4 heterodimer |
| $SMAD2_n$ | Nuclear SMAD2 |
| $pSMAD2_n$ | Nuclear phosphorylated SMAD2 |
| $SMAD2_n\text{-}SMAD4_n$ | Nuclear SMAD2–SMAD4 heterodimer |
| $pSMAD2_n\text{-}SMAD4_n$ | Nuclear pSMAD2–SMAD4 heterodimer |
| $SMAD3_c$ | Cytoplasmic SMAD3 |
| $pSMAD3_c$ | Cytoplasmic phosphorylated SMAD3 |
| $pSMAD3_{c,\mathrm{inh}}$ | Inhibited cytoplasmic SMAD3 |
| $SMAD3_c\text{-}ALK5_c\text{-}R2_c\text{-}TGF\beta_{\mathrm{act}}$ | Cytoplasmic TGF$\beta$ receptor complex with SMAD3 and active TGF-$\beta$ ligand |
| $pSMAD3_c\text{-}SMAD4_c$ | Cytoplasmic pSMAD3–SMAD4 heterodimer |
| $SMAD3_n$ | Nuclear SMAD3 |
| $pSMAD3_n$ | Nuclear phosphorylated SMAD3 |
| $SMAD3_n\text{-}SMAD4_n$ | Nuclear SMAD3–SMAD4 heterodimer |
| $pSMAD3_n\text{-}SMAD4_n$ | Nuclear pSMAD3–SMAD4 heterodimer |
| $SMAD4_c$ | Cytoplasmic SMAD4 |
| $SMAD4_n$ | Nuclear SMAD4 |

**Table S1** (***cont.***)

| Species | Description |
|---|---|
| $TGF\beta$ | Latent (inactive) TGF-$\beta$ |
| $TGF\beta_{\mathrm{act}}$ | Active TGF-$\beta$ |

#### Reactions for the three sub-models of the SB aortic calcification model

**Table S2:** ***Inflammation model***

| Reaction | Description |
|---|---|
| $\emptyset \xrightarrow{k_{\mathrm{inf1}}} LDL$ | LDL penetrates the endothelium and enters the leaflets |
| $LDL \xrightarrow{k_{\mathrm{inf2}}} \emptyset$ | LDL exits the leaflets through the endothelium |
| $LDL \xrightarrow{k_{\mathrm{inf3}}} oxLDL$ | LDL oxidation |
| $oxLDL + Macrophage \xrightarrow{k_{\mathrm{inf4}}} FoamCell$ | Macrophages absorb oxLDL to become lipid-laden foam cells |
| $oxLDL \xrightarrow{k_{\mathrm{inf5}}} oxLDL + Monocyte$ | Oxidized LDL promotes monocyte capture by increasing endothelial adhesion molecule expression, enabling circulating monocytes to enter the valve leaflets |
| $Monocyte \xrightarrow{k_{\mathrm{inf6}}} Macrophage$ | Monocytes differentiate into macrophages |
| $Monocyte \xrightarrow{k_{\mathrm{inf7}}} \emptyset$ | Monocyte apoptosis |
| $Macrophage + TGF\beta \xrightarrow{k_{\mathrm{inf8}}} Macrophage + TGF\beta_{\mathrm{act}}$ | Macrophages activate latent TGF-$\beta$ |
| $FoamCell + TGF\beta \xrightarrow{k_{\mathrm{inf8}}} FoamCell + TGF\beta_{\mathrm{act}}$ | Foam cells activate latent TGF-$\beta$ |
| $FoamCell \xrightarrow{k_{\mathrm{inf9}}} \emptyset$ | Foam cell apoptosis |
| $\emptyset \xrightarrow{k_{\mathrm{inf10}}} TGF\beta$ | TGF-$\beta$ secretion by macrophages and foam cells; the rate saturates at high cell numbers |
| $TGF\beta_{\mathrm{act}} \xrightarrow{k_{\mathrm{inf11}}} \emptyset$ | Activated TGF-$\beta$ degrades |
| $TGF\beta \xrightarrow{k_{\mathrm{inf12}}} \emptyset$ | Latent TGF-$\beta$ degrades |
| $Macrophage \xrightarrow{k_{\mathrm{inf13}}} \emptyset$ | Macrophage apoptosis |

**Table S3:** ***SMAD signaling model***

| Reaction | Description |
|---|---|
| $TGF\beta_{\mathrm{act}} + R2_s \underset{k_{\mathrm{sma1b}}}{\overset{k_{\mathrm{sma1a}}}{\rightleftharpoons}} R2_s\text{-}TGF\beta_{\mathrm{act}}$ | Activated TGF-$\beta$ binds and unbinds from TGF$\beta$RII at the VIC surface |
| $R2_s\text{-}TGF\beta_{\mathrm{act}} + ALK5_s \underset{k_{\mathrm{sma2b}}}{\overset{k_{\mathrm{sma2a}}}{\rightleftharpoons}} ALK5_s\text{-}R2_s\text{-}TGF\beta_{\mathrm{act}}$ | The TGF$\beta$–TGF$\beta$RII complex binds and unbinds from ALK5 (TGF$\beta$RI) |
| $ALK5_s\text{-}R2_s\text{-}TGF\beta_{\mathrm{act}} \overset{k_{\mathrm{sma3}}}{\rightleftharpoons} ALK5_c\text{-}R2_c\text{-}TGF\beta_{\mathrm{act}}$ | The receptor complex moves between the cell surface and cytoplasm |
| $ALK5_c\text{-}R2_c\text{-}TGF\beta_{\mathrm{act}} + SMAD2_c \underset{k_{\mathrm{sma4b}}}{\overset{k_{\mathrm{sma4a}}}{\rightleftharpoons}} SMAD2_c\text{-}ALK5_c\text{-}R2_c\text{-}TGF\beta_{\mathrm{act}}$ | The receptor complex binds and unbinds from SMAD2 |
| $ALK5_c\text{-}R2_c\text{-}TGF\beta_{\mathrm{act}} + SMAD3_c \underset{k_{\mathrm{sma4b}}}{\overset{k_{\mathrm{sma4a}}}{\rightleftharpoons}} SMAD3_c\text{-}ALK5_c\text{-}R2_c\text{-}TGF\beta_{\mathrm{act}}$ | The receptor complex binds and unbinds from SMAD3 |
| $SMAD2_c\text{-}ALK5_c\text{-}R2_c\text{-}TGF\beta_{\mathrm{act}} \xrightarrow{k_{\mathrm{sma5}}} ALK5_c\text{-}R2_c\text{-}TGF\beta_{\mathrm{act}} + pSMAD2_c$ | Bound SMAD2 is phosphorylated by the receptor complex |

**Table S3** (*cont.*)

| Reaction | Description |
|---|---|
| $SMAD3_c\text{-}ALK5_c\text{-}R2_c\text{-}TGF\beta_{\text{act}} \xrightarrow{k_{\text{sma5}}} ALK5_c\text{-}R2_c\text{-}TGF\beta_{\text{act}} + pSMAD3_c$ | Bound SMAD3 is phosphorylated by the receptor complex |
| $pSMAD2_c + SMAD4_c \underset{k_{\text{sma6b}}}{\overset{k_{sma6a}}{\rightleftharpoons}} pSMAD2_c\text{-}SMAD4_c$ | pSMAD2 heterodimerizes with SMAD4 |
| $pSMAD3_c + SMAD4_c \underset{k_{\text{sma6b}}}{\overset{k_{sma6a}}{\rightleftharpoons}} pSMAD3_c\text{-}SMAD4_c$ | pSMAD3 heterodimerizes with SMAD4 |
| $pSMAD2_c\text{-}SMAD4_c \xrightarrow{k_{\text{sma7}}} pSMAD2_n\text{-}SMAD4_n$ | pSMAD2–SMAD4 heterodimer translocates to the nucleus |
| $pSMAD3_c\text{-}SMAD4_c \xrightarrow{k_{\text{sma7}}} pSMAD3_n\text{-}SMAD4_n$ | pSMAD3–SMMAD4 heterodimer translocates to the nucleus |
| $pSMAD2_n\text{-}SMAD4_n \xrightarrow{k_{\text{sma8}}} SMAD2_n\text{-}SMAD4_n$ | Nuclear pSMAD2–SMAD4 is dephosphorylated |
| $pSMAD3_n\text{-}SMAD4_n \xrightarrow{k_{\text{sma8}}} SMAD3_n\text{-}SMAD4_n$ | Nuclear pSMAD3–SMAD4 is dephosphorylated |
| $SMAD2_n\text{-}SMAD4_n \xrightarrow{k_{\text{sma9}}} SMAD2_n + SMAD4_n$ | Nuclear SMAD2–SMAD4 heterodimer dissociates |
| $SMAD3_n\text{-}SMAD4_n \xrightarrow{k_{\text{sma9}}} SMAD3_n + SMAD4_n$ | Nuclear SMAD3–SMAD4 heterodimer dissociates |
| $SMAD2_c \underset{k_{\text{sma10b}}}{\overset{k_{\text{sma10a}}}{\rightleftharpoons}} SMAD2_n$ | SMAD2 shuttles between cytoplasm and nucleus |
| $SMAD3_c \underset{k_{\text{sma10b}}}{\overset{k_{\text{sma10a}}}{\rightleftharpoons}} SMAD3_n$ | SMAD3 shuttles between cytoplasm and nucleus |
| $SMAD4_c \underset{k_{\text{sma11b}}}{\overset{k_{\text{sma11a}}}{\rightleftharpoons}} SMAD4_n$ | SMAD4 shuttles between cytoplasm and nucleus |
| $\emptyset \underset{k_{\text{sma12b}}}{\overset{k_{\text{sma12a}}}{\rightleftharpoons}} R2_s$ | TGF$\beta$RII is generated and degraded |
| $\emptyset \underset{k_{\text{sma13b}}}{\overset{k_{\text{sma13a}}}{\rightleftharpoons}} ALK5_s$ | ALK5 (TGF$\beta$RI) is generated and degraded |
| $\emptyset \underset{k_{\text{sma14b}}}{\overset{k_{\text{sma14a}}}{\rightleftharpoons}} SMAD2_c$ | SMAD2 is generated and degraded |
| $\emptyset \underset{k_{\text{sma14b}}}{\overset{k_{\text{sma14a}}}{\rightleftharpoons}} SMAD3_c$ | SMAD3 is generated and degraded |
| $\emptyset \underset{k_{\text{sma15b}}}{\overset{k_{\text{sma15a}}}{\rightleftharpoons}} SMAD4_c$ | SMAD4 is generated and degraded |
| $ALK5_s\text{-}R2_s\text{-}TGF\beta_{\text{act}} \xrightarrow{k_{\text{sma16}}} \emptyset$ | The surface receptor complex is degraded |
| $pSMAD2_c \xrightarrow{k_{\text{sma17}}} pSMAD2_n$ | pSMAD2 translocates to the nucleus |
| $pSMAD3_c \xrightarrow{k_{\text{sma17}}} pSMAD3_n$ | pSMAD3 translocates to the nucleus |
| $pSMAD2_n + SMAD4_n \underset{k_{\text{sma18b}}}{\overset{k_{\text{sma18a}}}{\rightleftharpoons}} pSMAD2_n\text{-}SMAD4_n$ | Nuclear pSMAD2 binds and unbinds from SMAD4 |
| $pSMAD3_n + SMAD4_n \underset{k_{\text{sma18b}}}{\overset{k_{\text{sma18a}}}{\rightleftharpoons}} pSMAD3_n\text{-}SMAD4_n$ | Nuclear pSMAD3 binds and unbinds from SMAD4 |
| $pSMAD2_n \xrightarrow{k_{\text{sma19}}} SMAD2_n$ | Nuclear pSMAD2 is dephosphorylated |
| $pSMAD3_n \xrightarrow{k_{\text{sma19}}} SMAD3_n$ | Nuclear pSMAD3 is dephosphorylated |
| $pSMAD2_n \xrightarrow{k_{\text{sma20}}} \emptyset$ | Nuclear pSMAD2 is degraded |
| $pSMAD3_n \xrightarrow{k_{\text{sma20}}} \emptyset$ | Nuclear pSMAD3 is degraded |
| $R2_s \underset{k_{\text{sma21b}}}{\overset{k_{\text{sma21a}}}{\rightleftharpoons}} R2_c$ | TGF$\beta$RII moves between cytoplasm and surface |
| $ALK5_s \underset{k_{\text{sma22b}}}{\overset{k_{\text{sma22a}}}{\rightleftharpoons}} ALK5_c$ | ALK5 moves between cytoplasm and surface |

**Table S3 (*cont.*)**

| Reaction | Description |
|---|---|
| $ALK5_c\text{-}R2_c\text{-}TGF\beta_{\text{act}} \xrightarrow{k_{\text{sma23}}} ALK5_s\text{-}R2_s\text{-}TGF\beta_{\text{act}}$ | The receptor complex returns from the cytoplasm to the surface |
| $PKG_{\text{act}} + SMAD3_c \xrightarrow{k_{\text{sma24}}} PKG_{\text{act}} + pSMAD3_{c,\text{inh}}$ | Active PKG inhibits SMAD3 by phosphorylating inhibitory sites that prevent SMAD4 binding |
| $pSMAD3_{c,\text{inh}} \xrightarrow{k_{\text{sma25}}} SMAD3_c$ | Inhibited SMAD3 is restored to its active form |
| $pSMAD3_n\text{-}SMAD4_n \xrightarrow{k_{\text{sma26}}} pSMAD3_n\text{-}SMAD4_n + Ca$ | Nuclear pSMAD3–SMAD4 heterodimer promotes calcium production via transcription of pro-calcific genes |
| $pSMAD2_n\text{-}SMAD4_n \xrightarrow{k_{\text{sma27}}} pSMAD2_n\text{-}SMAD4_n + Ca$ | Nuclear pSMAD2–SMAD4 heterodimer promotes calcium production via transcription of pro-calcific genes |

**Table S4: *NO regulation model***

| Reaction | Description |
|---|---|
| $\emptyset \xrightarrow{k_{\text{NO}}} NO$ | NO production, with a shear-dependent rate constant |
| $NO + sGC \underset{k_{\text{inh1b}}}{\overset{k_{\text{inh1a}}}{\rightleftharpoons}} NO\text{-}sGC$ | NO binds and unbinds from sGC |
| $sGC + GTP \underset{k_{\text{inh2b}}}{\overset{k_{\text{inh2a}}}{\rightleftharpoons}} sGC\text{-}GTP$ | sGC binds and unbinds from GTP |
| $NO\text{-}sGC + GTP \underset{k_{\text{inh3b}}}{\overset{k_{\text{inh3a}}}{\rightleftharpoons}} NO\text{-}sGC\text{-}GTP$ | NO–sGC complex binds and unbinds from GTP |
| $sGC\text{-}GTP \xrightarrow{k_{\text{inh4}}} sGC + cGMP$ | sGC catalyzes conversion of GTP to cGMP |
| $NO\text{-}sGC\text{-}GTP \xrightarrow{k_{\text{inh5}}} NO\text{-}sGC + cGMP$ | NO–sGC catalyzes conversion of GTP to cGMP |
| $cGMP + PDE \underset{k_{\text{inh6b}}}{\overset{k_{\text{inh6a}}}{\rightleftharpoons}} PDE\text{-}cGMP$ | cGMP binds and unbinds from PDE |
| $PDE\text{-}cGMP \xrightarrow{k_{\text{inh7}}} PDE + GMP$ | PDE hydrolyzes cGMP to GMP |
| $NO \xrightarrow{k_{\text{inh8}}} \emptyset$ | NO degrades |
| $cGMP + PKG \xrightarrow{k_{\text{inh9}}} cGMP + PKG_{\text{act}}$ | cGMP activates PKG |
| $PKG_{\text{act}} \xrightarrow{k_{\text{inh10}}} PKG$ | Active PKG deactivates |
| $GMP \xrightarrow{k_{\text{inh11}}} GTP$ | GMP recycles back to GTP |

#### ODEs derived from the reactions in Tables S2–S4

$$[\dot{ALK5_c}] = \underbrace{k_{\text{sma22a}}[ALK5_s]}_{\text{ALK5 receptor goes into cytoplasm}} - \underbrace{k_{\text{sma22b}}[ALK5_c]}_{\text{ALK5 receptor reattaches to surface}} \tag{S1}$$

$$\begin{aligned}[ALK5_c\text{-}\dot{R2_c}\text{-}TGF\beta_{\text{act}}] = {} & \underbrace{k_{\text{sma5}}([SMAD2_c\text{-}ALK5_c\text{-}R2_c\text{-}TGF\beta_{\text{act}}] + [SMAD3_c\text{-}ALK5_c\text{-}R2_c\text{-}TGF\beta_{\text{act}}])}_{\text{Phosphorylation of SMAD2/3}} \\ & + \underbrace{k_{\text{sma3}}[ALK5_s\text{-}R2_s\text{-}TGF\beta_{\text{act}}]}_{\text{Receptor complex detaches from cell surface into cytoplasm}} \\ & + \underbrace{k_{\text{sma4b}}([SMAD2_c\text{-}ALK5_c\text{-}R2_c\text{-}TGF\beta_{\text{act}}] + [SMAD3_c\text{-}ALK5_c\text{-}R2_c\text{-}TGF\beta_{\text{act}}])}_{\text{Receptor complex dissociates from SMAD2/3}} \\ & - \underbrace{k_{\text{sma4a}}[ALK5_c\text{-}R2_c\text{-}TGF\beta_{\text{act}}]([SMAD2_c] + [SMAD3_c])}_{\text{Receptor complex binds to SMAD2/3}} - \underbrace{k_{\text{sma23}}[ALK5_c\text{-}R2_c\text{-}TGF\beta_{\text{act}}]}_{\text{Receptor complex breaks completely}}\end{aligned} \tag{S2}$$

$$\dot{[ALK5_s]} = \underbrace{k_{\text{sma22b}}[ALK5_c]}_{\text{ALK5 receptor reattaches to surface}} + \underbrace{k_{\text{sma13a}}}_{\text{ALK5 receptor synthesis}} + \underbrace{k_{\text{sma2b}}[ALK5_s\text{-}R2_s\text{-}TGF\beta_{\text{act}}]}_{\text{ALK5 dissociates from receptor complex}} - \underbrace{k_{\text{sma13b}}[ALK5_s]}_{\text{ALK5 receptor degradation}} - \underbrace{k_{\text{sma2a}}[ALK5_s][R2_s\text{-}TGF\beta_{\text{act}}]}_{\text{Receptor complex formation with ALK5}} - \underbrace{k_{\text{sma22a}}[ALK5_s]}_{\text{ALK5 receptor goes into cytoplasm}} \tag{S3}$$

$$\dot{[ALK5_s\text{-}R2_s\text{-}TGF\beta_{\text{act}}]} = \underbrace{k_{\text{sma2a}}[ALK5_s][R2_s\text{-}TGF\beta_{\text{act}}]}_{\text{Receptor complex formation with ALK5}} - \underbrace{k_{\text{sma2b}}[ALK5_s\text{-}R2_s\text{-}TGF\beta_{\text{act}}]}_{\text{ALK5 dissociates from receptor complex}} - \underbrace{k_{\text{sma3}}[ALK5_s\text{-}R2_s\text{-}TGF\beta_{\text{act}}]}_{\text{Receptor complex detaches from cell surface into cytoplasm}} - \underbrace{(k_{\text{sma16ld}} + k_{\text{sma16cd}})[ALK5_s\text{-}R2_s\text{-}TGF\beta_{\text{act}}]}_{\text{Ligand induced and constitutive degradation of receptor complex}} \tag{S4}$$

$$\dot{[Ca]} = \underbrace{k_{\text{sma26}}[pSMAD3_n\text{-}SMAD4_n]}_{\text{pSMAD3–4 transcription of calcific genes}} + \underbrace{k_{\text{sma27}}[pSMAD2_n\text{-}SMAD4_n]}_{\text{pSMAD2–4 transcription of calcific genes}} \tag{S5}$$

$$\dot{[FoamCell]} = \underbrace{k_{\text{inf4}}[oxLDL][Macrophage]}_{\text{Foam cell formation}} - \underbrace{k_{\text{inf9}}[FoamCell]}_{\text{Foam cell apoptosis}} \tag{S6}$$

$$\dot{[sGC]} = \underbrace{(k_{\text{inh4}} + k_{\text{inh2b}})[sGC\text{-}GTP]}_{\text{GTP unbinds or cGMP made}} + \underbrace{k_{\text{inh1b}}[NO\text{-}sGC]}_{\text{NO unbinds from sGC}} - \underbrace{k_{\text{inh2a}}[sGC][GTP]}_{\text{GTP binds sGC}} - \underbrace{k_{\text{inh1a}}[NO][sGC]}_{\text{NO binds sGC}} \tag{S7}$$

$$\dot{[sGC\text{-}GTP]} = \underbrace{k_{\text{inh2a}}[sGC][GTP]}_{\text{GTP binds sGC}} - \underbrace{(k_{\text{inh4}} + k_{\text{inh2b}})[sGC\text{-}GTP]}_{\text{cGMP made or GTP unbinds}} \tag{S8}$$

$$\dot{[GMP]} = \underbrace{k_{\text{inh7}}[PDE\text{-}cGMP]}_{\text{GMP made from cGMP–PDE complex}} - \underbrace{k_{\text{inh11}}[GMP]}_{\text{GMP recycled back to GTP}} \tag{S9}$$

$$\dot{[cGMP]} = \underbrace{k_{\text{inh5}}[NO\text{-}sGC\text{-}GTP]}_{\text{cGMP made from NO–sGC–GTP}} + \underbrace{k_{\text{inh4}}[sGC\text{-}GTP]}_{\text{cGMP made from sGC–GTP}} + \underbrace{k_{\text{inh6b}}[PDE\text{-}cGMP]}_{\text{cGMP unbinds from PDE}} - \underbrace{k_{\text{inh6a}}[cGMP][PDE]}_{\text{cGMP binds PDE}} \tag{S10}$$

$$\dot{[GTP]} = \underbrace{k_{\text{inh3b}}[NO\text{-}sGC\text{-}GTP]}_{\text{GTP unbinds from NO–sGC}} + \underbrace{k_{\text{inh2b}}[sGC\text{-}GTP]}_{\text{GTP unbinds from sGC}} + \underbrace{k_{\text{kinh11}}[GMP]}_{\text{GMP recycled back to GTP}} - \underbrace{k_{\text{inh2a}}[sGC][GTP]}_{\text{GTP binds sGC}} - \underbrace{k_{\text{inh3a}}[GTP][NO\text{-}sGC]}_{\text{GTP binds NO–sGC}} \tag{S11}$$

$$\dot{[LDL]} = \underbrace{k_{\text{inf1}}(\bar{\tau})}_{\text{Subendothelial LDL penetration}} - \underbrace{k_{\text{inf2}}[LDL]}_{\text{LDL diffusing out}} - \underbrace{k_{\text{inf3}}[LDL]}_{\text{LDL oxidation}} \tag{S12}$$

$$\dot{[oxLDL]} = \underbrace{k_{\text{inf3}}[LDL]}_{\text{LDL oxidation}} - \underbrace{k_{\text{inf4}}[oxLDL][Macrophage]}_{\text{Foam cell formation}} \tag{S13}$$

$$\dot{[Macrophage]} = \underbrace{k_{\text{inf6}}[Monocyte]}_{\text{Monocyte differentiation to macrophage}} - \underbrace{k_{\text{inf4}}[oxLDL][Macrophage]}_{\text{Foam cell formation}} - \underbrace{k_{\text{inf13}}[Macrophage]}_{\text{Macrophage apoptosis}} \tag{S14}$$

$$\dot{[Monocyte]} = \underbrace{k_{\text{inf5}}(\bar{\tau})[oxLDL]}_{\text{Baseline monocyte capture}} - \underbrace{k_{\text{inf6}}[Monocyte]}_{\text{Monocyte differentiation to macrophage}} - \underbrace{k_{\text{inf7}}[Monocyte]}_{\text{Monocyte apoptosis}} \tag{S15}$$

$$\dot{[NO]} = \underbrace{k_{\text{NO}}(\tau)}_{\text{Flow-induced NO production}} + \underbrace{k_{\text{inh1b}}[NO\text{-}sGC]}_{\text{NO unbinds from sGC}} - \underbrace{k_{\text{inh8}}[NO]}_{\text{NO degraded}} - \underbrace{k_{\text{inh1a}}[NO][sGC]}_{\text{NO binds to sGC}} \tag{S16}$$

$$\dot{[NO\text{-}sGC]} = \underbrace{k_{\text{inh1a}}[NO][sGC]}_{\text{NO binds sGC}} + \underbrace{(k_{\text{inh3b}} + k_{\text{inh5}})[NO\text{-}sGC\text{-}GTP]}_{\text{GTP unbinds or cGMP made}} - \underbrace{k_{\text{inh1b}}[NO\text{-}sGC]}_{\text{NO unbinds from sGC}} - \underbrace{k_{\text{inh3a}}[GTP][NO\text{-}sGC]}_{\text{GTP binds NO–sGC complex}} \tag{S17}$$

$$\dot{[NO\text{-}sGC\text{-}GTP]} = \underbrace{k_{\text{inh3a}}[GTP][NO\text{-}sGC]}_{\text{GTP binds NO–sGC complex}} - \underbrace{(k_{\text{inh3b}} + k_{\text{inh5}})[NO\text{-}sGC\text{-}GTP]}_{\text{GTP unbinds or cGMP made}} \tag{S18}$$

$$\dot{[PDE]} = \underbrace{(k_{\text{inh7}} + k_{\text{inh6b}})[PDE\text{-}cGMP]}_{\text{PDE released from cGMP–PDE}} - \underbrace{k_{\text{inh6a}}[cGMP][PDE]}_{\text{cGMP binds PDE}} \tag{S19}$$

$$\dot{[PDE\text{-}cGMP]} = \underbrace{k_{\text{inh6a}}[cGMP][PDE]}_{\text{cGMP binds PDE}} - \underbrace{(k_{\text{inh7}} + k_{\text{inh6b}})[PDE\text{-}cGMP]}_{\text{GMP made or cGMP unbinds}} \tag{S20}$$

$$\dot{[PKG]} = \underbrace{k_{\text{inh10}}[PKG_{\text{act}}]}_{\text{PKG autoinhibitory deactivation}} - \underbrace{k_{\text{inh9}}[PKG][cGMP]}_{\text{cGMP activates PKG}} \tag{S21}$$

$$\dot{[PKG_{\text{act}}]} = \underbrace{k_{\text{inh9}}[PKG][cGMP]}_{\text{cGMP activates PKG}} - \underbrace{k_{\text{inh10}}[PKG_{\text{act}}]}_{\text{PKG deactivates}} \tag{S22}$$

$$\dot{[R2_c]} = \underbrace{k_{\text{sma21a}}[R2_s]}_{\text{R2 receptor goes into cytoplasm}} - \underbrace{k_{\text{sma21b}}[R2_c]}_{\text{R2 receptor reattaches to surface}} \tag{S23}$$

$$\begin{aligned}\dot{[R2_s]} = {} & \underbrace{k_{\text{sma12a}}}_{\text{R2 receptor synthesis}} + \underbrace{k_{\text{sma23}}[ALK5_c\text{-}R2_c\text{-}TGF\beta_{\text{act}}]}_{\text{Receptor complex breaks completely}} + \underbrace{k_{\text{sma21b}}[R2_c]}_{\text{R2 receptor reattaches to surface}} \\ & + \underbrace{k_{\text{sma1b}}[R2_s\text{-}TGF\beta_{\text{act}}]}_{\text{TGF-}\beta\text{ unbinds from R2 receptor}} - \underbrace{k_{\text{sma12b}}[R2_s]}_{\text{R2 receptor degradation}} - \underbrace{k_{\text{sma1a}}[R2_s][TGF\beta_{\text{act}}]}_{\text{TGF-}\beta\text{ binds to R2 receptor}} - \underbrace{k_{\text{sma21a}}[R2_s]}_{\text{R2 receptor goes into cytoplasm}}\end{aligned} \tag{S24}$$

$$\begin{aligned}\dot{[R2_s\text{-}TGF\beta_{\text{act}}]} = {} & \underbrace{k_{\text{sma2b}}[ALK5_s\text{-}R2_s\text{-}TGF\beta_{\text{act}}]}_{\text{ALK5 dissociates from receptor complex}} + \underbrace{k_{\text{sma1a}}[R2_s][TGF\beta_{\text{act}}]}_{\text{TGF-}\beta\text{ binds to R2 receptor}} \\ & - \underbrace{k_{\text{sma2a}}[ALK5_s][R2_s\text{-}TGF\beta_{\text{act}}]}_{\text{Receptor complex formation with ALK5}} - \underbrace{k_{\text{sma1b}}[R2_s\text{-}TGF\beta_{\text{act}}]}_{\text{TGF-}\beta\text{ unbinds from R2 receptor}}\end{aligned} \tag{S25}$$

$$\begin{aligned}\dot{[SMAD2_c]} = {} & \underbrace{k_{\text{sma14a}}}_{\text{SMAD2/3 synthesis}} + \underbrace{k_{\text{sma4b}}[SMAD2_c\text{-}ALK5_c\text{-}R2_c\text{-}TGF\beta_{\text{act}}]}_{\text{Receptor complex dissociates from SMAD2/3}} + \underbrace{k_{\text{sma10b}}[SMAD2_n]}_{\text{Translocation out of nucleus}} \\ & - \underbrace{k_{\text{sma14b}}[SMAD2_c]}_{\text{SMAD2/3 degradation}} - \underbrace{k_{\text{sma4a}}[SMAD2_c][ALK5_c\text{-}R2_c\text{-}TGF\beta_{\text{act}}]}_{\text{Receptor complex binds to SMAD2/3}} - \underbrace{k_{\text{sma10a}}[SMAD2_c]}_{\text{Translocation into nucleus}}\end{aligned} \tag{S26}$$

$$\dot{[pSMAD2_c]} = \underbrace{k_{\text{sma5}}[SMAD2_c\text{-}ALK5_c\text{-}R2_c\text{-}TGF\beta_{\text{act}}]}_{\text{Receptor complex phosphorylates SMAD2/3}} + \underbrace{k_{\text{sma6b}}[pSMAD2_c\text{-}SMAD4_c]}_{\text{pSMAD2/3 unbinds from SMAD4}} - \underbrace{k_{\text{sma6a}}[SMAD4_c][pSMAD2_c]}_{\text{pSMAD2/3 binds to SMAD4 to form pSMAD2/3-4 complex}} - \underbrace{k_{\text{sma17}}[pSMAD2_c]}_{\text{pSMAD2/3 translocates into nucleus}} \tag{S27}$$

$$\dot{[SMAD2_c\text{-}ALK5_c\text{-}R2_c\text{-}TGF\beta_{\text{act}}]} = \underbrace{k_{\text{sma4a}}[SMAD2_c][ALK5_c\text{-}R2_c\text{-}TGF\beta_{\text{act}}]}_{\text{Receptor complex binds to SMAD2/3}} - \underbrace{(k_{\text{sma4b}} + k_{\text{sma5}})[SMAD2_c\text{-}ALK5_c\text{-}R2_c\text{-}TGF\beta_{\text{act}}]}_{\text{Dissociation and phosphorylation of SMAD2/3}} \tag{S28}$$

$$\dot{[pSMAD2_c\text{-}SMAD4_c]} = \underbrace{k_{\text{sma6a}}[SMAD4_c][pSMAD2_c]}_{\text{pSMAD2/3 binds to SMAD4}} - \underbrace{k_{\text{sma6b}}[pSMAD2_c\text{-}SMAD4_c]}_{\text{pSMAD2/3 unbinds from SMAD4}} - \underbrace{k_{\text{sma7}}[pSMAD2_c\text{-}SMAD4_c]}_{\text{pSMAD2/3-4 complex translocates into nucleus}} \tag{S29}$$

$$\dot{[SMAD2_n]} = \underbrace{k_{\text{sma19}}[pSMAD2_n]}_{\text{Dephosphorylation}} + \underbrace{k_{\text{sma9}}[SMAD2_n\text{-}SMAD4_n]}_{\text{Complex breaks}} + \underbrace{k_{\text{sma10a}}[SMAD2_c]}_{\text{Translocation into nucleus}} - \underbrace{k_{\text{sma10b}}[SMAD2_n]}_{\text{Translocation out of nucleus}} \tag{S30}$$

$$\dot{[pSMAD2_n]} = \underbrace{k_{\text{sma17}}[pSMAD2_c]}_{\text{Translocation into nucleus}} + \underbrace{k_{\text{sma18b}}[pSMAD2_n\text{-}SMAD4_n]}_{\text{Complex breaks}} - \underbrace{k_{\text{sma18a}}[pSMAD2_n][SMAD4_n]}_{\text{Binding to SMAD4}} - \underbrace{k_{\text{sma19}}[pSMAD2_n]}_{\text{Dephosphorylation}} - \underbrace{k_{\text{sma20}}[pSMAD2_n]}_{\text{Degradation}} \tag{S31}$$

$$\dot{[SMAD2_n\text{-}SMAD4_n]} = \underbrace{k_{\text{sma8}}[pSMAD2_n\text{-}SMAD4_n]}_{\text{Dephosphorylation}} - \underbrace{k_{\text{sma9}}[SMAD2_n\text{-}SMAD4_n]}_{\text{SMAD2/3-4 complex breaks}} \tag{S32}$$

$$\dot{[pSMAD2_n\text{-}SMAD4_n]} = \underbrace{k_{\text{sma18a}}[pSMAD2_n][SMAD4_n]}_{\text{Nuclear pSMAD2/3 binds to SMAD4}} + \underbrace{k_{\text{sma7}}[pSMAD2_c\text{-}SMAD4_c]}_{\text{Translocation into nucleus}} - \underbrace{k_{\text{sma8}}[pSMAD2_n\text{-}SMAD4_n]}_{\text{Dephosphorylation of pSMAD2/3-4 complex}} - \underbrace{k_{\text{sma18b}}[pSMAD2_n\text{-}SMAD4_n]}_{\text{Nuclear complex breaks}} \tag{S33}$$

$$\dot{[SMAD3_c]} = \underbrace{k_{\text{sma14a}}}_{\text{SMAD2/3 synthesis}} + \underbrace{k_{\text{sma4b}}[SMAD3_c\text{-}ALK5_c\text{-}R2_c\text{-}TGF\beta_{\text{act}}]}_{\text{Receptor complex dissociates from SMAD2/3}} + \underbrace{k_{\text{sma10b}}[SMAD3_n]}_{\text{Translocation out of nucleus}} + \underbrace{k_{\text{sma25}}[pSMAD3_{c,\text{inh}}]}_{\text{Cytosolic pSMAD3 desphosphorylates}} - \underbrace{k_{\text{sma14b}}[SMAD3_c]}_{\text{SMAD2/3 degradation}} - \underbrace{k_{\text{sma4a}}[SMAD3_c][ALK5_c\text{-}R2_c\text{-}TGF\beta_{\text{act}}]}_{\text{Receptor complex binds to SMAD2/3}} - \underbrace{k_{\text{sma24}}[PKG_{\text{act}}][SMAD3_c]}_{\text{PKG inhibits SMAD3 via hyperphosphorylation}} - \underbrace{k_{\text{sma10a}}[SMAD3_c]}_{\text{Translocation into nucleus}} \tag{S34}$$

$$\dot{[pSMAD3_c]} = \underbrace{k_{\text{sma5}}[SMAD3_c\text{-}ALK5_c\text{-}R2_c\text{-}TGF\beta_{\text{act}}]}_{\text{Receptor complex phosphorylates SMAD2/3}} + \underbrace{k_{\text{sma6b}}[pSMAD3_c\text{-}SMAD4_c]}_{\text{pSMAD2/3 unbinds from SMAD4}} - \underbrace{k_{\text{sma6a}}[SMAD4_c][pSMAD3_c]}_{\text{pSMAD2/3 binds to SMAD4 to form pSMAD2/3-4 complex}} - \underbrace{k_{\text{sma17}}[pSMAD3_c]}_{\text{pSMAD2/3 translocates into nucleus}} \tag{S35}$$

$$[p\dot{SMAD3}_{c,\mathrm{inh}}] = \underbrace{k_{\mathrm{sma24}}[PKG_{\mathrm{act}}][SMAD3_c]}_{\text{PKG inhibits SMAD3 via hyperphosphorylation}} - \underbrace{k_{\mathrm{sma25}}[pSMAD3_{c,\mathrm{inh}}]}_{\text{Cytosolic inhibited pSMAD3 desphosphorylates}} \tag{S36}$$

$$[SMAD3_c\text{-}A\dot{LK5}_c\text{-}R2_c\text{-}TGF\beta_{\mathrm{act}}] = \underbrace{k_{\mathrm{sma4a}}[SMAD3_c][ALK5_c\text{-}R2_c\text{-}TGF\beta_{\mathrm{act}}]}_{\text{Receptor complex binds to SMAD2/3}} - \underbrace{(k_{\mathrm{sma4b}} + k_{\mathrm{sma5}})[SMAD3_c\text{-}ALK5_c\text{-}R2_c\text{-}TGF\beta_{\mathrm{act}}]}_{\text{Dissociation and phosphorylation of SMAD2/3}} \tag{S37}$$

$$[pSMAD\dot{3}_c\text{-}SMAD4_c] = \underbrace{k_{\mathrm{sma6a}}[SMAD4_c][pSMAD3_c]}_{\text{pSMAD2/3 binds to SMAD4}} - \underbrace{k_{\mathrm{sma6b}}[pSMAD2_c\text{-}SMAD4_c]}_{\text{pSMAD2/3 unbinds from SMAD4}} - \underbrace{k_{\mathrm{sma7}}[pSMAD3_c\text{-}SMAD4_c]}_{\text{pSMAD2/3-4 complex translocates into nucleus}} \tag{S38}$$

$$[\dot{SMAD3}_n] = \underbrace{k_{\mathrm{sma19}}[pSMAD3_n]}_{\text{Dephosphorylation}} + \underbrace{k_{\mathrm{sma9}}[SMAD3_n\text{-}SMAD4_n]}_{\text{Complex breaks}} + \underbrace{k_{\mathrm{sma10a}}[SMAD3_c]}_{\text{Translocation into nucleus}} - \underbrace{k_{\mathrm{sma10b}}[SMAD3_n]}_{\text{Translocation out of nucleus}} \tag{S39}$$

$$[p\dot{SMAD3}_n] = \underbrace{k_{\mathrm{sma17}}[pSMAD3_c]}_{\text{Translocation into nucleus}} + \underbrace{k_{\mathrm{sma18b}}[pSMAD3_n\text{-}SMAD4_n]}_{\text{Complex breaks}} - \underbrace{k_{\mathrm{sma18a}}[pSMAD3_n][SMAD4_n]}_{\text{Binding to SMAD4}} - \underbrace{k_{\mathrm{sma19}}[pSMAD3_n]}_{\text{Dephosphorylation}} - \underbrace{k_{\mathrm{sma20}}[pSMAD3_n]}_{\text{Degradation}} \tag{S40}$$

$$[SMAD\dot{3}_n\text{-}SMAD4_n] = \underbrace{k_{\mathrm{sma8}}[pSMAD3_n\text{-}SMAD4_n]}_{\text{Dephosphorylation}} - \underbrace{k_{\mathrm{sma9}}[SMAD3_n\text{-}SMAD4_n]}_{\text{SMAD2/3-4 complex breaks}} \tag{S41}$$

$$[pSMAD\dot{3}_n\text{-}SMAD4_n] = \underbrace{k_{\mathrm{sma18a}}[pSMAD3_n][SMAD4_n]}_{\text{Nuclear pSMAD2/3 binds to SMAD4}} + \underbrace{k_{\mathrm{sma7}}[pSMAD3_c\text{-}SMAD4_c]}_{\text{Translocation into nucleus}} - \underbrace{k_{\mathrm{sma8}}[pSMAD3_n\text{-}SMAD4_n]}_{\text{Dephosphorylation of pSMAD2/3-4 complex}} - \underbrace{k_{\mathrm{sma18b}}[pSMAD3_n\text{-}SMAD4_n]}_{\text{Nuclear complex breaks}} \tag{S42}$$

$$[\dot{SMAD4}_c] = \underbrace{k_{\mathrm{sma15a}}}_{\text{SMAD4 synthesis}} + \underbrace{k_{\mathrm{sma6b}}([pSMAD2_c\text{-}SMAD4_c] + [pSMAD3_c\text{-}SMAD4_c])}_{\text{pSMAD2/3 unbinds from SMAD4}} + \underbrace{k_{\mathrm{sma11b}}[SMAD4_n]}_{\text{SMAD4 translocates out of nucleus}} - \underbrace{k_{\mathrm{sma15b}}[SMAD4_c]}_{\text{SMAD4 degradation}} - \underbrace{k_{\mathrm{sma6a}}[SMAD4_c]([pSMAD2_c] + [pSMAD3_c])}_{\text{pSMAD2/3 binds to SMAD4 to form pSMAD2/3-4 complex}} - \underbrace{k_{\mathrm{sma11a}}[SMAD4_c]}_{\text{SMAD4 translocates into nucleus}} \tag{S43}$$

$$[\dot{SMAD4}_n] = \underbrace{k_{\mathrm{sma9}}([SMAD2_n\text{-}SMAD4_n] + [SMAD3_n\text{-}SMAD4_n])}_{\text{Complex breaks}} + \underbrace{k_{\mathrm{sma18b}}([pSMAD2_n\text{-}SMAD4_n] + [pSMAD3_n\text{-}SMAD4_n])}_{\text{Nuclear complex breaks}} + \underbrace{k_{\mathrm{sma11a}}[SMAD4_c]}_{\text{SMAD4 translocates into nucleus}} - \underbrace{k_{\mathrm{sma11b}}[SMAD4_n]}_{\text{SMAD4 translocates out of nucleus}} - \underbrace{k_{\mathrm{sma18a}}([pSMAD2_n] + [pSMAD3_n])[SMAD4_n]}_{\text{Nuclear pSMAD2/3 binds to SMAD4}} \tag{S44}$$

$$[\dot{TGF\beta}] = \underbrace{k_{\text{inf10}}}_{\text{LatentTGF-}\beta\text{ production}} - \underbrace{k_{\text{inf12}}[TGF\beta]}_{\text{LatentTGF-}\beta\text{ degradation}} - \underbrace{k_{\text{inf8}}([FoamCell] + [Macrophage])[TGF\beta]}_{\text{TGF-}\beta\text{ activation}} \quad \text{(S45)}$$

$$[\dot{TGF\beta}_{\text{act}}] = \underbrace{k_{\text{inf8}}([FoamCell] + [Macrophage])[TGF\beta]}_{\text{TGF-}\beta\text{ activation}} + \underbrace{k_{\text{sma1b}}[R2_s\text{-}TGF\beta_{\text{act}}]}_{\text{TGF-}\beta\text{ unbinds from R2 receptor}}$$
$$+ \underbrace{k_{\text{sma23}}[ALK5_c\text{-}R2_c\text{-}TGF\beta_{\text{act}}]}_{\text{Receptor complex breaks completely}} - \underbrace{k_{\text{sma1a}}[R2_s][TGF\beta_{\text{act}}]}_{\text{TGF-}\beta\text{ binds to R2 receptor}} - \underbrace{k_{\text{inf11}}[TGF\beta_{\text{act}}]}_{\text{ActiveTGF-}\beta\text{ degradation}} \quad \text{(S46)}$$

#### Initial species concentrations and rate constants

**Table S5:** ***Initial species concentrations for the integrated SB model***

| Species | Initial Amount (M) | Module(s) | Reference |
|---|---|---|---|
| $ALK5_c$ | $1.19 \times 10^{-11}$ | SMAD signaling | Chung et al.[1] |
| $ALK5_s$ | $1.33 \times 10^{-12}$ | SMAD signaling | Chung et al.[1] |
| *FoamCell* | 0 | Inflammation | Arzani et al.[2] |
| *sGC* | $3.2 \times 10^{-10}$ | NO regulation | Garmaroudi et al.[3] |
| *cGMP* | 0 | NO regulation | Garmaroudi et al.[3] |
| *GTP* | $9.75 \times 10^{-7}$ | NO regulation | Garmaroudi et al.[3] |
| *LDL* | 0 | Inflammation | Arzani et al.[2] |
| *oxLDL* | 0 | Inflammation | Arzani et al.[2] |
| *Macrophage* | 0 | Inflammation | Arzani et al.[2] |
| *Monocyte* | 0 | Inflammation | Arzani et al.[2] |
| *NO* | $3.84 \times 10^{-11}$ | NO regulation | He and Liu[4] |
| *PDE* | $4.8 \times 10^{-10}$ | NO regulation | Garmaroudi et al.[3] |
| *PKG* | $3.2 \times 10^{-8}$ | NO regulation | Inferred from Keilbach et al.[5] |
| $PKG_{\text{act}}$ | 0 | SMAD signaling, NO regulation | N/A |
| $R2_c$ | $1.19 \times 10^{-11}$ | SMAD signaling | Chung et al.[1] |
| $R2_s$ | $1.33 \times 10^{-12}$ | SMAD signaling | Chung et al.[1] |
| $SMAD2_c$ | $2.26 \times 10^{-10}$ | SMAD signaling | Chung et al.[1] |
| $SMAD2_n$ | $3.98 \times 10^{-11}$ | SMAD signaling | Chung et al.[1] |
| $SMAD3_c$ | $2.26 \times 10^{-10}$ | SMAD signaling | Chung et al.[1] |
| $SMAD3_n$ | $3.98 \times 10^{-11}$ | SMAD signaling | Chung et al.[1] |
| $SMAD4_c$ | $3.31 \times 10^{-10}$ | SMAD signaling | Chung et al.[1] |
| $SMAD4_n$ | $3.45 \times 10^{-11}$ | SMAD signaling | Chung et al.[1] |
| $TGF\beta$ | 0 | Inflammation | Arzani et al.[2] |
| $TGF\beta_{\text{act}}$ | 0 | Inflammation, SMAD signaling | Arzani et al.[2] |

In Table S5, the initial concentrations of PKG and $\text{PKG}_{\text{act}}$ are based on estimates from Keilbach et al.,[5] who reported tissue concentrations for PKG of ~10 pmol/g of wet tissue. Assuming a tissue density of 1 g/mL, this corresponds to a tissue concentration on the order of $10^{-8}$ M. Because the reported values in the study were somewhat higher, we adopt this estimate as the total PKG concentration and set the initial concentration of PKG accordingly. The active form, $PKG_{\text{act}}$, is initialized at 0.

**Table S6:** ***Rate constants for the inflammation pathway model***

| Parameter | Value | Definition | Reference |
|---|---|---|---|
| $\bar{k}_{\text{inf1}}$ | $3.78 \times 10^{-6}$ M/s | Baseline LDL penetration | Arzani et al.[2] |
| $k_{\text{inf1}}$ | *See Eq.* (S47) | Subendothelial LDL penetration | Arzani et al.[2] |
| $k_{\text{inf2}}$ | $2.40 \times 10^{-5}$ s$^{-1}$ | LDL out diffusion | Arzani et al.[2] |
| $k_{\text{inf3}}$ | $3 \times 10^{-4}$ s$^{-1}$ | LDL oxidation | Arzani et al.[2] |
| $k_{\text{inf4}}$ | $1.71 \times 10^{-16}$ M$^{-1}$s$^{-1}$ | Foam cell formation | Arzani et al.[2] |
| $\bar{k}_{\text{inf5}}$ | $6.37 \times 10^{5}$ s$^{-1}$ | Baseline monocyte capture | Arzani et al.[2] |
| $k_{\text{inf5}}$ | *See Eq.* (S48) | Monocyte capture via LDL | Arzani et al.[2] |
| $k_{\text{inf6}}$ | $1.15 \times 10^{-6}$ s$^{-1}$ | Monocyte → macrophage differentiation | Arzani et al.[2] |
| $k_{\text{inf7}}$ | $2.76 \times 10^{-6}$ s$^{-1}$ | Monocyte apoptosis | Arzani et al.[2] |
| $k_{\text{inf8}}$ | $7.41 \times 10^{-13}$ M$^{-1}$s$^{-1}$ | TGF-$\beta$ activation | Arzani et al.[2] |
| $k_{\text{inf9}}$ | $1.22 \times 10^{-8}$ s$^{-1}$ | Foam cell apoptosis | Inferred from Jiang et al.[6] and Bajpai et al.[7] |
| $k_{\text{inf10}}$ | *See Eq.* (S49) | Latent TGF-$\beta$ production | Arzani et al.[2] |
| $k_{\text{inf11}}$ | $5.78 \times 10^{-3}$ s$^{-1}$ | Active TGF-$\beta$ degradation | Arzani et al.[2] |
| $k_{\text{inf12}}$ | $1.28 \times 10^{-4}$ s$^{-1}$ | Latent TGF-$\beta$ degradation | Arzani et al.[2] |
| $k_{\text{inf13}}$ | $1.32 \times 10^{-9}$ s$^{-1}$ | Macrophage apoptosis | Estimated from Bajpai et al.[7] |

In Table S6, $k_{\text{inf1}}$ is based on Arzani et al.,[2] who reported a baseline LDL penetration rate, which we denote as $\bar{k}_{\text{inf1}}$. This value is multiplied by a factor $(1+\bar{\tau}/\bar{\tau}_{\text{ref}})^{-1}$ to account for the effects of WSS on LDL penetration, i.e.,

$$k_{\text{inf1}}(\tau) = \frac{\bar{k}_{\text{inf1}}}{1+\bar{\tau}/\bar{\tau}_{\text{ref}}}. \tag{S47}$$

Similarly, $k_{\text{inf5}}$ is based on the second-order rate constant reported by Arzani et al.[2] for the interaction between monocytes and LDL. We multiply this by the same scaling factor to account for variations in WSS, i.e.,

$$k_{\text{inf5}}(\tau) = \frac{\bar{k}_{\text{inf5}}}{1+\bar{\tau}/\bar{\tau}_{\text{ref}}}. \tag{S48}$$

Additionally, we define the rate constant for foam cell apoptosis as $k_{\text{inf9}} = 9.28 \cdot k_{\text{inf13}}$, where $k_{\text{inf13}}$ is the macrophage apoptosis rate constant. This is based on results from Bajpai et al.,[7] who estimated the percentage of sex-mismatched CCR2$^{+}$ macrophages in the body after 8.8 years. They also reported that oxLDL-treated macrophages had a 47.7% apoptosis rate after 48h, compared to 5.14% in control, which is a ∼9.28-fold increase in the apoptosis rate for foam cells.

Finally, the rate constant for latent TGF-$\beta$ production, $k_{\text{inf10}}$, is assumed to depend on the total concentration of immune cells (*FoamCell* + *Macrophage*) and $V_{\text{valve}}$ (Table 1), which assumes different values depending on valve thickness,

$$k_{\text{inf10}} = \alpha \cdot \frac{3.3 \times 10^{-7} \cdot ([\mathit{FoamCell}] + [\mathit{Macrophage}]) \cdot V_{\text{valve}}}{\alpha \cdot ([\mathit{FoamCell}] + [\mathit{Macrophage}]) \cdot V_{\text{valve}} \cdot \psi + 2.84 \times 10^{4}}. \tag{S49}$$

Here, $\alpha$ is the foam cell formation ratio (See Table 1) and $\psi$ is a unit conversion factor defined as *Desired units* = *Current units*/$\psi$. In Arzani et al.,[2] immune cell concentrations are in g/m$^3$ and $V_{\text{valve}}$ is in m$^3$. This gives $k_{\text{inf10}}$ in units of g/(s·m$^3$). The conversion factor $\psi$ is used here to convert $k_{\text{inf10}}$ into units of M/s, consistent with the other sub-models in the SB model.

**Table S7:** ***Rate constants for the SMAD signaling pathway model***

| Parameter | Value | Description | Reference |
|---|---|---|---|
| $k_{\text{sma1a}}$ | $4.14 \times 10^{10}$ M$^{-1}$s$^{-1}$ | TGF-$\beta$ binds to R2 receptor | Chung et al.[1] |
| $k_{\text{sma1b}}$ | $4.97 \times 10^{-3}$ s$^{-1}$ | TGF-$\beta$ unbinds from R2 receptor | Chung et al.[1] |
| $k_{\text{sma2a}}$ | $4.14 \times 10^{10}$ M$^{-1}$s$^{-1}$ | Receptor complex formation with ALK5 | Chung et al.[1] |
| $k_{\text{sma2b}}$ | $4.97 \times 10^{-3}$ s$^{-1}$ | ALK5 dissociates from receptor complex | Chung et al.[1] |
| $k_{\text{sma3}}$ | $6.58 \times 10^{-3}$ s$^{-1}$ | Receptor complex detaches from the cell surface | Chung et al.[1] |
| $k_{\text{sma4a}}$ | $9.42 \times 10^{8}$ M$^{-1}$s$^{-1}$ | Receptor complex binds to SMAD2/3 | Chung et al.[1] |
| $k_{\text{sma4b}}$ | $1.62 \times 10^{-2}$ s$^{-1}$ | Receptor complex dissociates from SMAD2/3 | Chung et al.[1] |
| $k_{\text{sma5}}$ | $7.47 \times 10^{2}$ s$^{-1}$ | Receptor complex phosphorylates SMAD2/3 | Chung et al.[1] |
| $k_{\text{sma6a}}$ | $3.77 \times 10^{10}$ M$^{-1}$s$^{-1}$ | pSMAD2/3 binds to SMAD4 to form pSMAD2/3-4 | Chung et al.[1] |
| $k_{\text{sma6b}}$ | $2.43 \times 10^{1}$ s$^{-1}$ | pSMAD2/3 unbinds from SMAD4 | Chung et al.[1] |
| $k_{\text{sma7}}$ | $1.35 \times 10^{-2}$ s$^{-1}$ | pSMAD2/3-4 complex translocates into nucleus | Chung et al.[1] |
| $k_{\text{sma8}}$ | $4.2 \times 10^{-4}$ s$^{-1}$ | pSMAD2/3-4 complex dephosphorylates | Chung et al.[1] |
| $k_{\text{sma9}}$ | $1.68 \times 10^{-3}$ s$^{-1}$ | SMAD2/3-4 complex breaks | Chung et al.[1] |
| $k_{\text{sma10a}}$ | $2.7 \times 10^{-3}$ s$^{-1}$ | SMAD2/3 translocates into nucleus | Chung et al.[1] |
| $k_{\text{sma10b}}$ | $5.8 \times 10^{-3}$ s$^{-1}$ | SMAD2/3 translocates out of nucleus | Chung et al.[1] |
| $k_{\text{sma11a}}$ | $3.35 \times 10^{-4}$ s$^{-1}$ | SMAD4 translocates into nucleus | Chung et al.[1] |
| $k_{\text{sma11b}}$ | $2.9 \times 10^{-3}$ s$^{-1}$ | SMAD4 translocates out of nucleus | Chung et al.[1] |
| $k_{\text{sma12a}}$ | $3.54 \times 10^{-16}$ M/s | R2 receptor synthesis | Chung et al.[1] |
| $k_{\text{sma12b}}$ | $4.67 \times 10^{-4}$ s$^{-1}$ | R2 receptor degradation | Chung et al.[1] |
| $k_{\text{sma13a}}$ | $3.54 \times 10^{-16}$ M/s | ALK5 receptor synthesis | Chung et al.[1] |
| $k_{\text{sma13b}}$ | $4.67 \times 10^{-4}$ s$^{-1}$ | ALK5 receptor degradation | Chung et al.[1] |
| $k_{\text{sma14a}}$ | $1.21 \times 10^{-15}$ M/s | SMAD2/3 synthesis | Chung et al.[1] |
| $k_{\text{sma14b}}$ | $1.08 \times 10^{-5}$ s$^{-1}$ | SMAD2/3 degradation | Chung et al.[1] |
| $k_{\text{sma15a}}$ | $2.21 \times 10^{-15}$ M/s | SMAD4 synthesis | Chung et al.[1] |
| $k_{\text{sma15b}}$ | $2 \times 10^{-5}$ s$^{-1}$ | SMAD4 degradation | Chung et al.[1] |
| $k_{\text{sma16ld}}$ | $6.58 \times 10^{-3}$ s$^{-1}$ | Ligand induced degradation of receptor complex | Chung et al.[1] |
| $k_{\text{sma16cd}}$ | $4.67 \times 10^{-4}$ s$^{-1}$ | Constitutive degradation of receptor complex | Chung et al.[1] |
| $k_{\text{sma17}}$ | $8.38 \times 10^{-3}$ s$^{-1}$ | pSMAD2/3 translocates into nucleus | Chung et al.[1] |
| $k_{\text{sma18a}}$ | $1.05 \times 10^{9}$ M$^{-1}$s$^{-1}$ | Nuclear pSMAD2/3 binds to SMAD4 | Chung et al.[1] |
| $k_{\text{sma18b}}$ | $1.52 \times 10^{-2}$ s$^{-1}$ | Nuclear pSMAD2/3-4 complex breaks | Chung et al.[1] |
| $k_{\text{sma19}}$ | $4.2 \times 10^{-4}$ s$^{-1}$ | Nuclear pSMAD2/3 dephosphorylates | Chung et al.[1] |
| $k_{\text{sma20}}$ | $9 \times 10^{-5}$ s$^{-1}$ | Nuclear SMAD2/3 degradation | Chung et al.[1] |
| $k_{\text{sma21a}}$ | $6.58 \times 10^{-3}$ s$^{-1}$ | R2 receptor goes into cytoplasm | Chung et al.[1] |
| $k_{\text{sma21b}}$ | $6.58 \times 10^{-4}$ s$^{-1}$ | R2 receptor reattaches to surface | Chung et al.[1] |
| $k_{\text{sma22a}}$ | $6.58 \times 10^{-3}$ s$^{-1}$ | ALK5 receptor goes into cytoplasm | Chung et al.[1] |
| $k_{\text{sma22b}}$ | $6.58 \times 10^{-4}$ s$^{-1}$ | ALK5 receptor reattaches to surface | Chung et al.[1] |
| $k_{\text{sma23}}$ | $6.58 \times 10^{-4}$ s$^{-1}$ | Receptor complex breaks completely | Chung et al.[1] |
| $k_{\text{sma24}}$ | $5 \times 10^{3}$ M$^{-1}$s$^{-1}$ | PKG inhibits SMAD3 via hyperphosphorylation | Chung et al.[1] |
| $k_{\text{sma25}}$ | $4.20 \times 10^{-4}$ s$^{-1}$ | Cytosolic inhibited pSMAD3 desphosphorylates | Chung et al.[1] |
| $k_{\text{sma26}}$ | *See Eq.* (**??**) | pSMAD3-4 transcribes pro-calcific genes | N/A |
| $k_{\text{sma27}}$ | *See Eq.* (**??**) | pSMAD2-4 transcribes pro-calcific genes | N/A |

**Table S8:** ***Rate constants for the NO regulation model***

| Parameter | Value | Description | Reference |
|---|---|---|---|
| $k_{\text{NO}}$ | *See Fig. S1* | NO synthesis | Sriram et al.[8] |
| $k_{\text{inh1a}}$ | $0.78 \times 10^6$ $\text{M}^{-1}\text{s}^{-1}$ | NO binds to sGC | Inferred from Sharina and Martin[9] |
| $k_{\text{inh1b}}$ | $1.8 \times 10^{-1}$ $\text{s}^{-1}$ | NO unbinds from sGC | Sayed et al.[10] |
| $k_{\text{inh2a}}$ | $2.5 \times 10^5$ $\text{M}^{-1}\text{s}^{-1}$ | GTP binds to sGC | *See text below* |
| $k_{\text{inh2b}}$ | $1.8 \times 10^2$ $\text{s}^{-1}$ | GTP unbinds from sGC | Garmaroudi et al.[3] |
| $k_{\text{inh3a}}$ | $3.13 \times 10^7$ $\text{M}^{-1}\text{s}^{-1}$ | GTP binds to NO-sGC | *See text below* |
| $k_{\text{inh3b}}$ | $4.3 \times 10$ $\text{s}^{-1}$ | GTP unbinds from NO-sGC | Garmaroudi et al.[3] |
| $k_{\text{inh4}}$ | $1.5 \times 10^{-1}$ $\text{s}^{-1}$ | cGMP made from GTP-sGC | Garmaroudi et al.[3] |
| $k_{\text{inh5}}$ | $2.87 \times 10$ $\text{s}^{-1}$ | cGMP made from NO-GTP-sGC | Garmaroudi et al.[3] |
| $k_{\text{inh6a}}$ | $3.13 \times 10^8$ $\text{M}^{-1}\text{s}^{-1}$ | cGMP binds to PDE | Garmaroudi et al.[3] |
| $k_{\text{inh6b}}$ | $1.30 \times 10^{-1}$ $\text{s}^{-1}$ | cGMP unbinds from PDE | Garmaroudi et al.[3] |
| $k_{\text{inh7}}$ | $2.2$ $\text{s}^{-1}$ | GMP made from cGMP-PDE | Garmaroudi et al.[3] |
| $k_{\text{inh8}}$ | $1.4$ $\text{s}^{-1}$ | NO degradation | Thomas et al.[11] |
| $k_{\text{inh9}}$ | $3.13 \times 10^4$ $\text{M}^{-1}\text{s}^{-1}$ | cGMP activates PKG | *See text below* |
| $k_{\text{inh10}}$ | $5 \times 10^{-3}$ $\text{s}^{-1}$ | PKG autoinhibitory deactivation | Inferred from Srihirun et al.[12] |
| $k_{\text{inh11}}$ | $1.5 \times 10^{-3}$ $\text{s}^{-1}$ | GMP recycled back to GTP | Inferred from Khan et al.[13] and Mc-kee et al.[14] |

In Table S8, the rate constant for NO–sGC binding, $k_{\text{inh1a}}$ is based on the intrinsic cellular association rate constant $k_{\text{on}} = 1.4$–$4.5 \times 10^8$ $\text{M}^{-1}\text{s}^{-1}$, reported by Sharina et al.[9] Because sGC-expressing cells occupy less volume than the full leaflet, by roughly two orders of magnitude, most NO molecules diffuse without encountering the enzyme. Following the homogenization framework of Nicholson and Bassingthwaighte,[15,16] we scale $k_{\text{on}}$ by this volumetric factor to obtain an effective tissue-level rate constant $k_{\text{eff}} \approx 0.45$–$1.44 \times 10^6$ $\text{M}^{-1}\text{s}^{-1}$. Based on this, we set $k_{\text{inh1a}} = 0.78 \times 10^6$ $\text{M}^{-1}\text{s}^{-1}$.

For $k_{\text{inh3a}}$, the rate constant for GTP binding to the NO–sGC complex, we adopted a tissue-level rate corresponding to a cellular value of $\sim 10^5$ $\text{M}^{-1}\text{s}^{-1}$. The use of a lower effective rate is intentional: $k_{\text{inh3a}}$ is treated as a phenomenological parameter that captures system-level constraints, such as spatial hindrance, allosteric effects, or post-translational modifications, which may reduce the efficiency of GTP association with NO-bound sGC *in vivo*. A similar rationale is used for $k_{\text{inh2a}}$, which describes GTP binding to inactive sGC. Here, we set $k_{\text{inh2a}} < k_{\text{inh3a}}$, consistent with reports that NO binding induces a conformational change that increases the affinity of sGC for GTP. By encoding this mechanistic asymmetry, rather than assuming equal binding rates,[3] the model more faithfully represents the sequential activation and catalytic behavior of sGC.

For $k_{\text{inh9}}$, although cGMP binds PKG rapidly *in vitro*, we again employ a lower effective rate constant to represent macroscopic factors such as intracellular buffering, compartmentalization, and cooperative activation. The calibrated value yields physiologically reasonable PKG dynamics. For the PKG deactivation rate constant $k_{\text{inh10}}$, we selected a conservative value of $5 \times 10^{-3}$ $\text{s}^{-1}$ based on work by Srihirun et al.,[12] who reported a peak in the PKG activity marker P-VASP Ser239 at approximately 15 min, follwed by a return to baseline by 60 min. Fitting an exponential decay to a 5% residual yields $t_{1/2} \approx 10.4$ min ($k \approx 1.1 \times 10^{-3}$ $\text{s}^{-1}$). However, since the signal may have fully decayed, or simply fallen below the $\sim$ 5% noise floor, we adopt $k = 5 \times 10^{-3}$ $\text{s}^{-1}$ to ensure complete deactivation within the observed experimental window.

Finally, the rate constant for GMP $\rightarrow$ GTP recycling, $k_{\text{inh11}}$, was chosen to lie between experimentally grounded lower and upper bounds. A lower bound of $\sim 7.7 \times 10^{-5}$ $\text{s}^{-1}$ was inferred from McKee et al.,[14] who observed 13% guanine nucleotide conversion over 30 min in isolated cardiac mitochondria. As an upper bound, we computed an effective first-order rate constant, $k_{\text{inh11}}^{\text{upper}} = k_{\text{cat}}[E]/K_M$, from Michaelis–Menten kinetics (assuming $[\text{GMP}] \ll K_M$) using parameters from Figure 6E of Khan et al.[13] and an enzyme concentration of 10 nM, matching their assays. The resulting values spanned 0.015–0.045 $\text{s}^{-1}$ across six variants. Because this study characterized only GMP $\rightarrow$ GDP conversion and the assumption $[\text{GMP}] \ll K_M$ may fail as GMP accumulates, the true physiological rate likely falls below this range. Thus, our choice of $1.5\times10^{-3}$ s$^{-1}$ represents a conservative, physiologically plausible estimate.

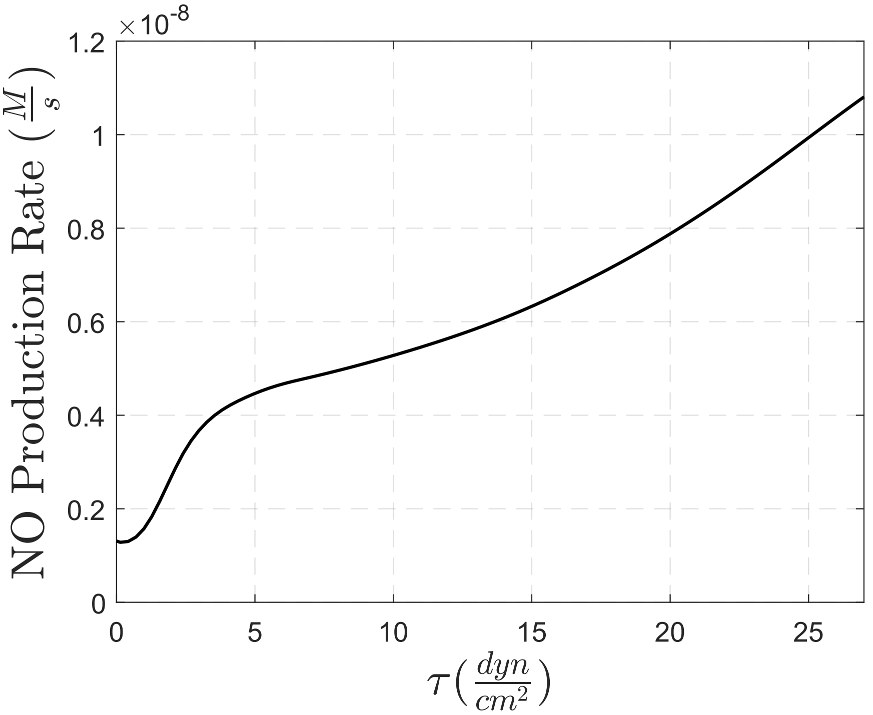


**Figure S1:** *NO production rate (in M/s) vs. shear stress (in dynes per* cm$^2$*). Taken from Sriram et al.*[8]

#### Sensitivity analysis

##### Method

To quantify the influence of individual model parameters on long-term calcification outcomes, we performed a local, one-at-a-time sensitivity analysis. For a generic model parameter $\theta$, sensitivity was assessed by symmetrically perturbing $\theta$ about its nominal value while holding all other parameters fixed. Specifically, parameter values were perturbed as $\theta(1 \pm s)$, and the resulting changes in calcification were evaluated at the terminal simulation time of 23 years.

The sensitivity coefficient was defined as

$$S_\theta = \frac{\mathrm{Ca}(t = 23 \text{ years}; \theta(1+s)) - \mathrm{Ca}(t = 23 \text{ years}; \theta(1-s))}{2\,\mathrm{Ca}(t = 23 \text{ years}; \theta)}, \tag{S50}$$

where $\mathrm{Ca}(t;\theta)$ denotes the model-predicted calcification level at time $t$ for parameter value $\theta$. This normalized metric represents the average relative change in terminal calcification induced by a symmetric perturbation of the parameter and enables direct comparison of sensitivities across parameters with differing units and magnitudes.

All biochemical and kinetic model parameters were perturbed using $s = 0.10$ ($\pm 10\%$), consistent with standard local sensitivity analysis practice. For biomechanical inputs, specifically wall shear stress and strain, larger perturbations of $s = 0.15$ ($\pm 15\%$) were employed to account for intrinsic cycle-to-cycle variability in these quantities arising from the coupled flow–structure interaction dynamics. Sensitivities were computed independently for each leaflet thickness configuration and reported using the absolute value of $S_\theta$ to rank parameter influence on calcification progression.

##### Stress and strain input sensitivity analysis

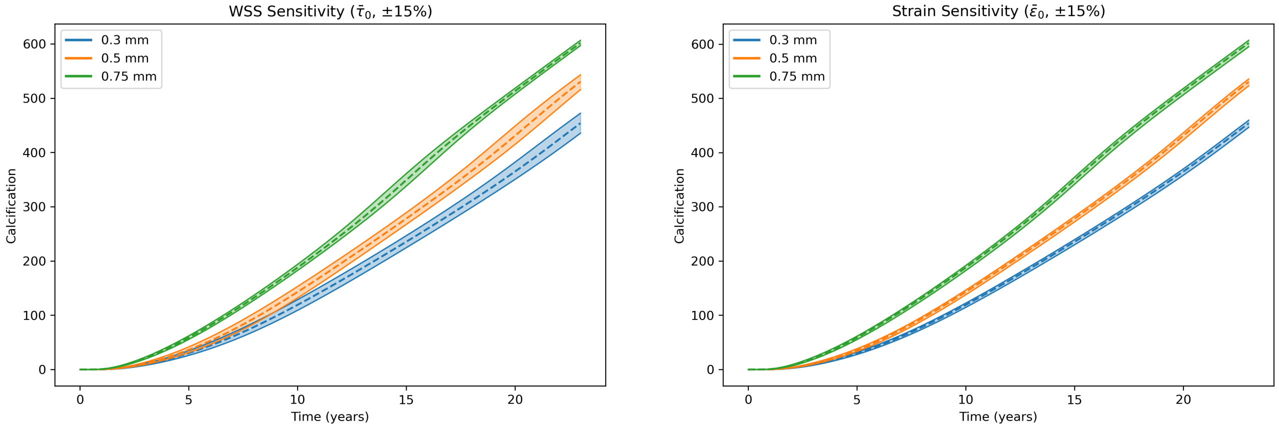


**Figure S2:** ***Local sensitivity envelopes for biomechanical inputs.*** *Shown are the model-predicted calcification trajectories resulting from symmetric ±15% perturbations to the wall shear stress (left) and strain (right) inputs for all three leaflet thickness cases. Shaded regions denote the envelope bounded by the upper and lower perturbation trajectories, while dashed curves indicate the corresponding baseline simulations. Larger perturbations were applied to these biomechanical inputs to account for intrinsic cycle-to-cycle variability arising from the coupled flow–structure interaction dynamics.*

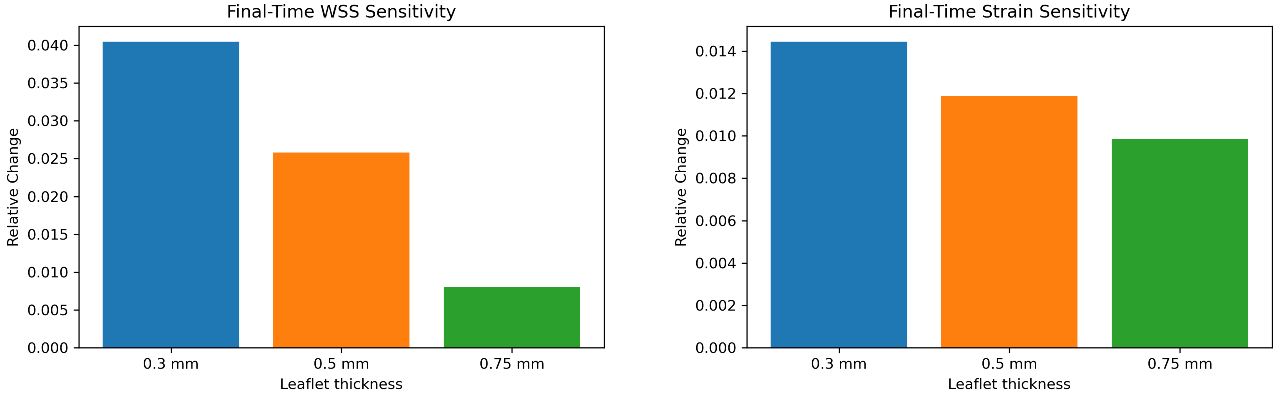


**Figure S3:** ***Sensitivities of calcification to biomechanical inputs across leaflet thicknesses.*** *Bars show the relative change in terminal calcification at* $t = 23$ *years resulting from symmetric perturbations to wall shear stress (left) and strain (right) for each leaflet thickness case. Sensitivities are computed independently for each input and thickness configuration and quantify the dependence of long-term calcification outcomes on biomechanical loading conditions.*

#### Model parameter sensitivity analysis

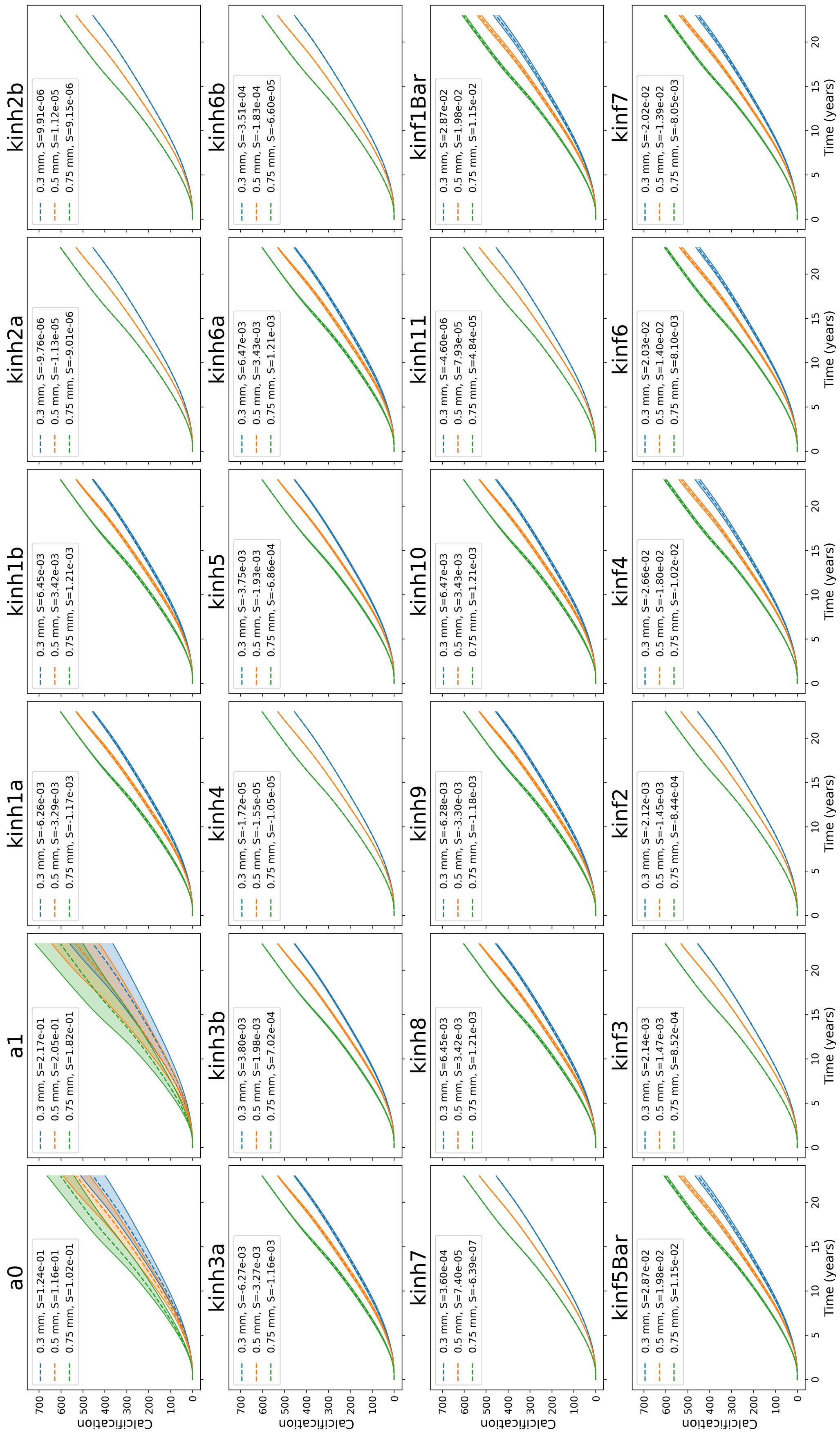


**Figure S4:** ***Local sensitivity envelopes for all model parameters across leaflet thicknesses.*** *Each panel shows the calcification response to symmetric* $\pm 10\%$ *perturbations for a single parameter, with shaded envelopes corresponding to the three leaflet thickness cases. The dotted curve is the baseline simulation.*

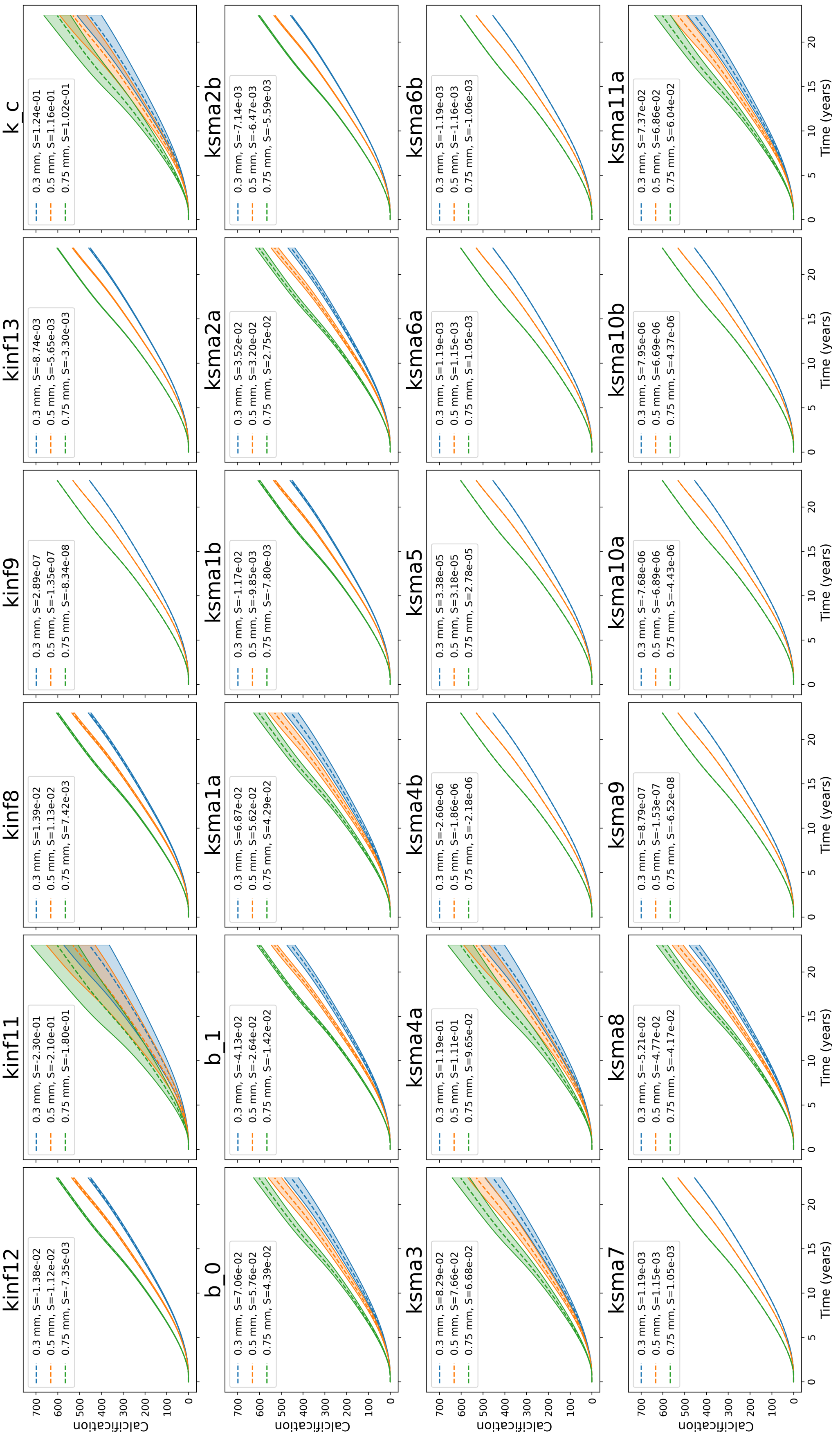


**Figure S4:** *(continued)*

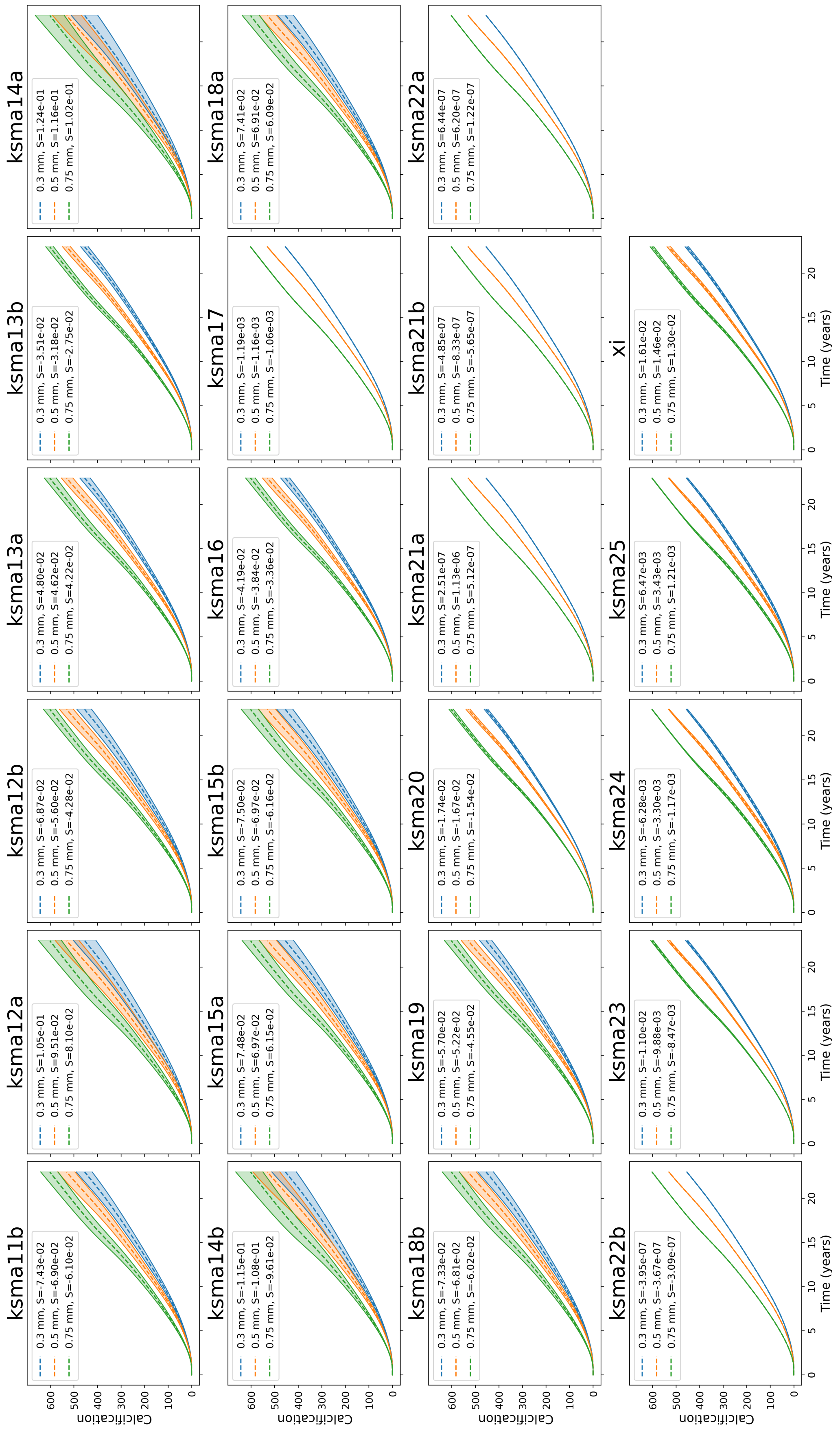


**Figure S4:** *(continued)*

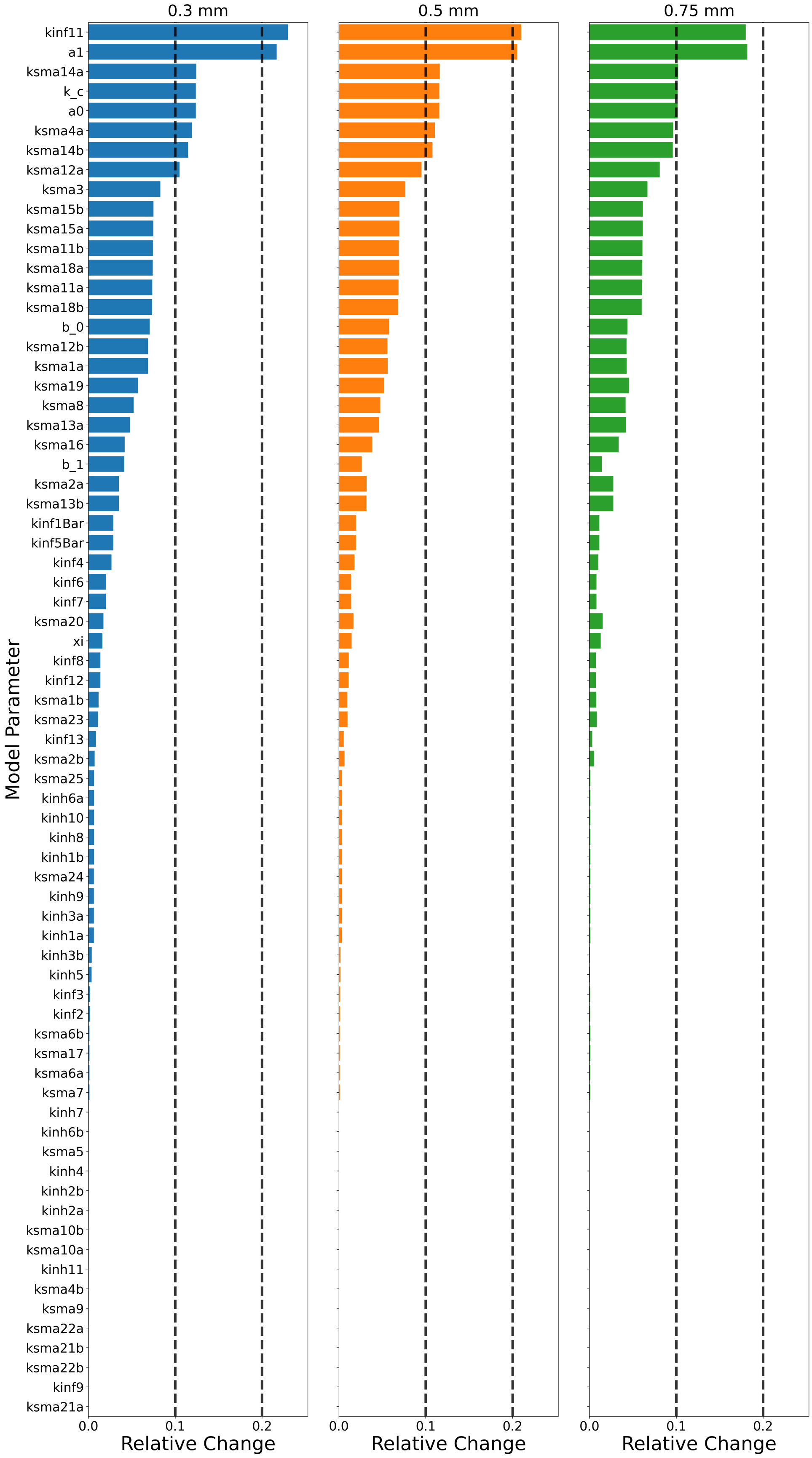


**Figure S5:** ***Ranked sensitivity of terminal calcification to model parameters.*** *Horizontal bar plots show the relative sensitivity of terminal calcification to individual model parameters for each leaflet thickness case. Parameters are ranked according to sensitivity for the 0.3 mm leaflet, and the same ordering is used for the thicker leaflet cases to facilitate direct comparison. Sensitivity values quantify the relative influence of each parameter on long-term calcification outcomes.*

[4] Guo-Wei He and Zhi-Gang Liu. Comparison of nitric oxide release and endothelium-derived hyperpolarizing factor–mediated hyperpolarization between human radial and internal mammary arteries. *Circulation*, 104(suppl_1):I–344, 2001.
